# Multi-omics analysis identifies LBX1 and NHLH1 as central regulators of human midbrain dopaminergic neuron differentiation

**DOI:** 10.1101/2023.01.27.525898

**Authors:** Borja Gomez Ramos, Jochen Ohnmacht, Nikola de Lange, Aurélien Ginolhac, Elena Valceschini, Aleksandar Rakovic, Rashi Halder, François Massart, Christine Klein, Roland Krause, Marcel H. Schulz, Thomas Sauter, Rejko Krüger, Lasse Sinkkonen

## Abstract

Midbrain dopaminergic neurons (mDANs) control voluntary movement, cognition, and reward behavior under physiological conditions and are implicated in human diseases such as Parkinson’s disease (PD). Many transcription factors (TFs) controlling human mDAN differentiation during development have been described, but much of the regulatory landscape remains undefined. Using a tyrosine hydroxylase (TH) iPSC reporter line, we have generated time series transcriptomic and epigenomic profiles of purified mDANs during differentiation. Integrative analysis predicted novel central regulators of mDAN differentiation and super-enhancers were used to prioritize key TFs. We find LBX1, NHLH1 and NR2F1/2 to be necessary for mDAN differentiation and show that overexpression of either LBX1 or NHLH1 can also improve mDAN specification. NHLH1 is necessary for the induction of neuronal miR-124, while LBX1 regulates cholesterol biosynthesis, possibly through mTOR signaling. Consistently, rapamycin treatment led to an inhibition of mDAN differentiation. Thus, our work reveals novel regulators of human mDAN differentiation.

## Introduction

Induced pluripotent stem cell (iPSC) technology presents a unique system to study how transcription factors (TFs) control cell differentiation and specification in humans. TFs bind to genomic regulatory regions such as enhancers and promoters to mediate their action. The TFs occupying an enhancer region collectively control transcriptional initiation of their target genes through different mechanisms (1, 2). In particular super-enhancers (SEs), dense clusters of enhancers under high regulatory load (3), have been associated with cell identity genes, master TFs, and are enriched with disease-associated genetic variants (4, 5). Identifying TF binding profiles and active enhancer regions across the genome requires laborious and indirect methods such as chromatin immunoprecipitation sequencing (ChIP-seq). The presence of regulatory proteins alone does not necessarily indicate active gene regulation. To overcome this limitation, these methods can be combined with gene expression analysis in an integrative multiomics approach to determine the role of specific TFs or enhancers in gene regulation. Moreover, a time course analysis combining transcriptomic and epigenomic data has shown great potential for studying processes like cell differentiation and revealing the regulatory events’ hierarchy. Such approaches have been able to generate gene regulatory networks (GRN) that also take into account the regulatory landscape of the cells, facilitating the identification of key TFs (6–8).

Midbrain dopaminergic neurons (mDANs) are widely used in biomedical research due to their involvement in different psychiatric and neurological disorders such as schizophrenia, drug addiction and PD (9–11). The current protocols for generating mDANs from iPSCs produce heterogeneous populations and incompletely specified cells (12, 13). The genetic program underlying mDAN development has been extensively investigated (reviewed in 14), with most of the studies relying on transcriptomic data and mice as the model organism. With the emergence of single-cell technologies, improved insights into human and mouse midbrain development have revealed differences in the temporal dynamics, cell composition, and expression of TFs (13, 15).

All these findings highlight our limited understanding of human development, hampering our ability to apply the developmental knowledge to improve iPSC differentiation protocols. Only a few studies have considered the epigenetic landscape of mDANs during development (16–18). Furthermore, cellular heterogeneity present in the iPSC-derived cultures obscure physiological or biological insights from the cell type of interest (19, 20).

In this study, we profiled differentiating human iPSC-derived mDANs at the transcriptomic and epigenomic levels. Integrative analysis using our EPIC-DREM pipeline (8, 21–23) generated timepoint-specific gene regulatory interactions. Together with mapping of cell type-specific SEs, this allowed the identification of putative key TFs controlling mDANs. We show that LBX1, NHLH1 and NR2F1/2 are necessary for mDAN differentiation, with LBX1 and NHLH1 also able to increase the number of mDANs. Further characterization of these TFs revealed the control of cholesterol biosynthesis by LBX1 and induction of miR-124 by NHLH1 as a few of the mechanisms contributing to mDAN specification. In summary, this study provides novel profiling of differentiating mDANs. Our data can be exploited for further purposes such as studies on disease-associated regulatory genetic variation.

## Materials and Methods

### Cell lines

The human iPSC line GM17602 (Coriell) was used in this study as a control and for the generation of a tyrosine hydroxylase (TH) reporter cell line. This iPSC line was previously characterized in (24) (called HFF) and used in (19) for the generation of the reporter line. Briefly, in the reporter line, the T2A coding sequence was fused with the mCherry open reading frame, and it was biallelically inserted in place of the stop codon in the endogenous TH locus using CRISPR/Cas9 editing (Figure 1A).

**Figure 1:**
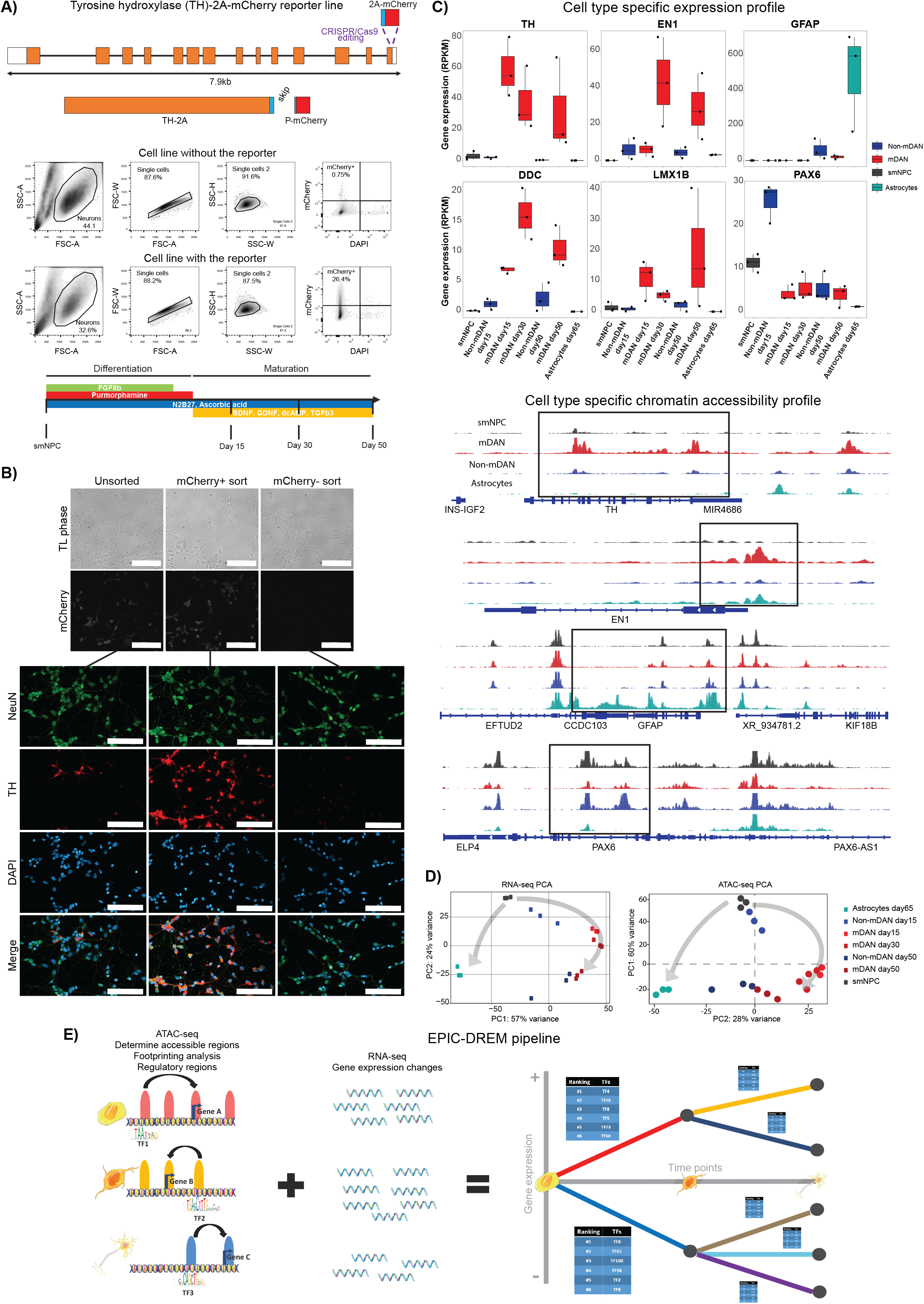
Epigenomic and transcriptomic analysis of human mDAN differentiation. **A)** Gene editing scheme of the human iPSC reporter line, a representative flow cytometry comparison of iPSC-derived neurons from the lines with and without the reporter, and the differentiation protocol used with the different time points analyzed. **B)** Validation of the reporter line by immunocytochemistry. TH staining correlated with mCherry signal. NeuN was used as a neuronal marker while DAPI stained all nuclei. Scale bar = 100 µm. **C)** Gene expression and chromatin accessibility profiles from the different samples analyzed showed specific enrichments for cell type specific markers of mDANs (TH, EN1, DDC, LMX1B), astrocytes (GFAP), and non-mDANs (PAX6). ATAC-seq tracks are plotted under the same scale for comparison purposes. N = 3 for all datasets and samples. **D)** PCA plots of the RNA-seq and ATAC-seq data reveal the differentiation dynamics. **E)** EPIC-DREM pipeline scheme showing the different steps in data integration for the generation of time point-specific gene regulatory networks across differentiation. ATAC-seq was used to define gene regulatory regions and the TFs associated to them were identified through footprinting analysis. Integration with the transcriptional changes defined by RNA-seq allowed the creation of sets of co-expressed genes associated with a ranking of TFs controlling them across time.

### Cell culture and differentiation

hiPSC maintenance, generation of small molecule neural precursor cells (smNPC) and differentiation towards mDANs are described in (25). In Figure 1A, a schematic representation of the mDAN differentiation protocol can be observed with the different media used and for how long the cells were kept in culture. This differentiation protocol is composed of three different media. smNPCs are incubated in Differentiation medium one containing 100 ng/ml of FGF8b, 1 µM of purmorphamine (PMA), and 200 µM of ascorbic acid which starts with smNPC and is used during the first eight days of differentiation. From day 8 until day 10 of differentiation cells are kept in Differentiation medium two, composed of 0.5 µM PMA and 200 µM of ascorbic acid. Lastly, maturation medium containing 200 µM of ascorbic acid, 10 ng/ml of brain-derived neurotrophic factor (BDNF), 10 ng/ml of glial cell-derived neurotrophic factor (GDNF), 500 µM of dcAMP, and 1 ng/ml of TGFβ3 which is used from day 10 until the desired time point. The molecules used in the three different media were mixed in N2B27 medium (1:1 of Dulbecco’s modified Eagle medium/Nutrient Mixture F-12 [DMEM/F12] and Neurobasal medium supplemented with 1% Pen/Strep, 1% GlutaMAX, 1% B27 supplement minus vitamin A, and 0.5% N2 supplement, all from Gibco).

Astrocyte differentiation was induced based on the protocol from (26), with small changes. Briefly, smNPC were seeded in N2B27 medium complemented with 3 µM CHIR99021, 0.5 µM PMA and 150 µM ascorbic acid. After two days, fresh medium plus 20 ng/ml of FGF-2 was added to the smNPC culture. On day four, the cells were split into a new plate to start neural stem cell (NSC) generation. The medium used for the generation and maintenance of NSC contained DMEM/F-12, 1% Pen/Strep, 1% GlutaMAX, 1% B27 supplement serum-free (with vitamin A), 1% N2 supplement, 40 ng/ml EGF, 40n g/ml FGF-2, and 1.5 ng/ul hLIF. The cells were kept in this medium for 3-4 passages. Then, astrocyte differentiation was started in DMEM/F-12 supplemented with 1% Pen/Strep, 1% GlutaMAX, and 1% FBS.

All the plates used were previously coated with Geltrex matrix from Gibco.

### Flow cytometry and FACS

On the day of analysis, medium was removed from the cells and Accutase (Gibco) was added to the well. Cells were incubated in Accutase at 37°C until detachment (10-30 min). Then, 2 volumes of DMEM/F-12 were added to the well. Cells were gently pipetted up and down until dissociation and collected in a 15 ml Falcon tube after passing them through a 50 µm cell strainer to obtain a single-cell suspension. Falcon tubes were centrifuged 3 min at 300xg and room temperature (RT). Pellets were resuspended in PBS and cells were transferred to a 1.5 ml Eppendorf tube. 4’,6-diamidino-2-phenylindole (DAPI) was added to the cells at 5 µg/ml and incubated for 5 min at 4°C in a tube rotator. After incubation, the cells were washed twice with 2% (w/v) BSA in PBS. Then, cells were ready for either flow cytometry analysis or fluorescence-assisted cell sorting (FACS). For flow cytometry analysis, the BD LSRFortessa™ cell analyzer was used. For FACS, the BD FACSAria™ III sorter was used. FlowJo version 10 software was used for data processing and generation of plots.

### Total RNA extraction, cDNA synthesis and RT-qPCR

For total RNA extraction, Quick-RNA miniprep kit from Zymo Research (R1055) was used as per manufacturer’s instructions. When total RNA had to be extracted from FACS sorted cells, Quick-RNA microprep kit from Zymo Research (R1050) was used. Briefly, to avoid RNA degradation due to FACS, cells were collected in batches for no longer than 10 min. Then, sorted cells were pelleted by centrifugation for 3min at 500 g and 4°C. Lysis buffer from the kit was added to the pellet until 150 000 – 300 000 cells were collected. RNA extraction was finalized following manufacturer’s instruction.

For cDNA synthesis, amounts from 200 ng to 1 ug of total RNA were used, depending on sample availability. To perform the reaction, dNTPs (0.5 mM, ThermoFisher, R0181), oligo dT-primer (2.5 µM), 1µl RevertAid reverse transcriptase (200 U/µl, ThermoFisher, EP0441), and 1µl Ribolock RNase inhibitor (40 U/µl, ThermoFisher, EO0381) were mixed with the proper amount of total RNA in a final volume of 40 µl. cDNA synthesis was performed at 42°C for 1 hr. Reaction was terminated by incubating the reaction at 70°C for 10 min. cDNA was diluted 1:10, 1:5 or 1:2 in DNase/RNase free water, depending on the amount of total RNA used (1 µg, 500 ng or 200 ng, respectively). Diluted cDNA was stored at −20°C.

RT-qPCR was performed in an Applied Biosystems 7500 Fast Real-Time PCR system. For the reaction, 5 µl of diluted cDNA was mixed with 1xAbsolute Blue qPCR SYBR green low ROX mix (ThermoFisher, AB4322B) and 500 nM primer concentration, in a final volume of 20 µl per well using AmpliStart 96well plate (Westburg, WB1900). PCR reaction had the following settings: 95°C for 15min and then 40 cycles of 95°C for 15sec, 55°C for 15sec and 72°C for 30sec. The 2^-(ΔΔCt)^ method was used to calculate gene expression levels. ΔΔCt was calculated using the following formula: (ΔCt_(target gene)_-ΔCt_(housekeeping gene)_) - (ΔCt_(target gene)_-ΔCt_(housekeeping gene)_)_reference condition_. ACTB was used as the housekeeping gene. GraphPad Prism 9 was used to create plots and perform statistics. Data was always normalized to the reference condition and one sample t test was performed to determine significance.

### Omni-ATAC

The Omni-ATAC protocol was performed with 50 000 cells (or 25 000 cells for day 50 neurons) with minor modifications from (27). Using TDE1 Tagment DNA Enzyme and TD Buffer from Illumina. Cells were either derived from FACS or directly from cell culture plates after detachment with Accutase. After amplification with primer sets as described in (28), libraries underwent size selection using AMPure XP beads (Beckman Coulter) to remove large fragments. Libraries were stored at −20°C. For library quality control, the Agilent High Sensitivity DNA kit (5067-4626) was used in a 2100 Bioanalyzer instrument. Libraries used for sequencing presented a fragment distribution starting from ∼200 bp until around 1000 bp, with nucleosomal pattern peaks.

### Low-input ChIP-seq and calling of SEs

To perform low-input chromatin immunoprecipitation (ChIP) for H3K27ac, the Low Cell ChIP-Seq Kit from Active Motif was used (53084). For smNPC and sorted neurons, a total of 150 000 and 200 000 cells were used, respectively. The protocol was performed according to manufacturer’s instructions, with minor changes. Sonication of the cells was performed using a Bioruptor® Pico sonication device from Diagenode. The settings used for sonication were 40 cycles of 30 sec off and 30 sec on at 8°C. After sonication, 20% of the sonicated sample volume was saved as input. For the immunoprecipitation (IP) reaction, a total of 4 µg of H3K27ac antibody was used per reaction. Details about the antibody can be found in the Antibodies section. Input samples that were collected after sonication were processed together with IP reactions starting from the step where reversal of cross-links and DNA purification was performed. After this point, samples were processed in parallel. Libraries were stored at −20°C until quality control and sequencing. For library quality control, the same procedure as in the Omni-ATAC protocol was used. For low-input ChIP, good libraries presented an average of 600 bp fragment distribution. The low-input ChIP-seq protocol was validated by comparing the identified SEs to those previously detected in smNPC via regular high-input ChIP-seq methods (29). SEs were considered as regions larger than 10 kb.

### Immunocytochemistry

PhenoPlate 96-well (PerkinElmer, 6055308) previously coated with geltrex was used for immunocytochemistry. Cells were fixed in 4% paraformaldehyde for 15 min at RT. Then, cells were washed three times with PBS. For permeabilization, cells were incubated 1 hr at RT in PBS, 0.4% Triton-X, 10% goat serum and 2% BSA. After 1 hr, cells were washed twice with PBS. Primary antibody was diluted in PBS, 0.1% Triton-X, 1% goat serum and 0.2% BSA. Cells were incubated with primary antibody shaking overnight at 4°C. Next day, cells were washed three times with PBS. Then, secondary antibody was diluted in the same buffer as the primary and incubated for 2-3 hrs at RT. Finally, cells were washed three times with PBS. In the first wash, DAPI was added to the PBS (5 µg/ml) and incubated for 15 min at RT. Images were taken using a Zeiss spinning disk confocal microscope. Image processing was done in ImageJ.

### Bacterial culture, plasmid extraction and lentivirus production

Glycerol stocks of bacteria containing the plasmid of interest were taken from −80°C. Without letting the glycerol stocks thaw and with the help of a P10 pipette tip, bacteria were added to a polypropylene graduated culture tube (Roth, EC01.1) containing 5 ml of LB Broth medium (20.6 g/l, Roth, X968.2) supplemented with Ampicillin (100 µg/ml). Bacteria were incubated at 37°C and 120 rpm shaking for 5 hrs. Then, bacteria were transferred to an Erlenmeyer containing 150 ml of LB Broth medium supplemented with ampicillin. Erlenmeyer was incubated overnight at 37°C and shaken at 120 rpm. After bacterial expansion, plasmid extraction was performed using NucleoBond Xtra Midi EF (740420.50) as per manufacturer’s instructions.

For lentivirus production, 8 million HEK293T cells were seeded in a T75 flask using 15 ml of DMEM (Gibco) supplemented with 1% Pen/Strep, and 10% FBS and transfected the next day for lentiviral production. Third-generation lentiviral particles were produced. Briefly, 4 µg of pMDG, 2 µg of pMDL, 2 µg of pREV, and 8 µg of the plasmid of interest were mixed with 200 µl of CaCl2 (1 M, Sigma, 21115-100ML). Volume was completed with sterile water up to 800 µl. The 800 µl of the plasmid mixture was mixed with 800 µl of HEPES buffered saline (Sigma, 51558-50ML) by making bubbles slowly in a dropwise manner. This transfection mixture was incubated for 20 min at RT. In the meanwhile, 16 µl of 25mM chloroquine was added to the T75 flask containing the HEK293T cells and incubated for a minimum of 5 min to facilitate transfection. Next, the transfection mixture was added to the HEK293T cells. After 4-6hr, medium was removed from HEK293T cells and 14 ml of fresh medium was added. After 48 hrs, lentiviral particles were ready for collection. HEK293T medium from the flask was collected in a 15ml Falcon tube and centrifuged for 10 min at 2000 rpm and 4°C. The supernatant was cleared by filtering through a 0.45 µm filter (Sartorious, 16537). Filtered lentiviral particles were aliquoted in cryovials (1 ml aliquots) and stored at −80°C.

### Transduction

Two different approaches for transduction of differentiating neurons were used in this study: early and late transductions. All the lentiviral particles used contained a GFP reporter that helped control for transduction efficiency. For overexpression constructs a codon-optimized cDNA sequence of the TF was used. Lentiviral particles were previously tested to adjust transduction efficiencies to ∼80%, determined by GFP positive cells using flow cytometry.

For early transduction, smNPC were seeded in a 6-well plate with a density of 1-2 million cells per well and differentiation was started by seeding them directly in a differentiation medium. Next day, medium was removed and lentiviral particles were added to the cells in a final volume of 1 ml. Plate was sealed with parafilm and centrifuged for 10 min at 250 g and RT for spinfection. After spinfection, 1 ml of differentiation medium was added to the cells. Lentiviral particles were incubated overnight. Next day, lentiviral particles were removed, and fresh differentiation medium was added to the cells. Differentiation continued until the day of analysis.

For late transduction, differentiating cells were split on day 8 of differentiation into a 6 well plate at a density of 3 million cells per well. The next day, the medium was removed and transduction was performed as described for early transduction. Differentiation continued until the day of analysis.

### GW3965, ABX464, and Rapamycin treatments

GW3965-HCl (Selleckchem, S2630), ABX464 (Selleckchem, S0076), and Rapamycin (Selleckchem, S1039) molecules were tested to determine their working concentration in our cultures. For GW3965-HCl, 0.1 µM was sufficient to induce SREBF1 mRNA levels. For ABX464, 10 µM was sufficient to induce miR-124-3p. For Rapamycin, 10 nM was sufficient to downregulate SREBF1 mRNA levels.

The effect of the molecules was tested during normal differentiation and under LBX1 or NHLH1 knockdown (KD) conditions. Briefly, neuronal differentiations were started and on day 8 of differentiation, cells were split into a 6-well plate with a density of 3 million cells per well. Next day, late transduction with shLBX1, shNHLH1 or shScramble was performed as described above. On day 10 of differentiation, transduced cultures were treated either with GW3965, ABX464, or rapamycin. Cells transduced with shLBX1 were treated with GW3965 or Rapamycin, while cells transduced with shNHLH1 were treated with ABX464. Treatments continued until day 15, when cells were analyzed.

### TaqMan assay

TaqMan assay was performed to determine miR-124-3p levels using TaqMan™ MicroRNA Reverse Transcription Kit (ThermoFisher, 4366596), TaqMan™ MicroRNA Assay hsa-miR-124-3p (ThermoFisher, AssayID 003188_mat, 4440886), TaqMan™ MicroRNA Control Assay U6 snRNA (ThermoFisher, AssayID 001973, 4427975), TaqMan™ MicroRNA Assay hsa-miR-423-5p (ThermoFisher, AssayID 002340, 4427975), and TaqMan™ Fast Advanced Master Mix (ThermoFisher, 4444556). First, reverse transcription reaction was performed using 0.15 µl of 100 mM dNTPs, 1 µl of MultiScribe™ Reverse Transcriptase (50 U/µl), 1.5 µl of 10X Reverse Transcription buffer, 0.19 µl of RNase inhibitor (20 U/µl), 10 ng of total RNA, and 3 µl of RT primer (either U6 or miR-124-3p primer) in a final volume of 15 µl. Reactions were incubated for 30 min at 16°C and then 30 min at 42°C. Reaction was terminated by incubating the tubes for 5 min at 85°C. Then, PCR was performed using the same plates and machine as for RT-qPCR. For the PCR reaction, 1 µl of 20X TaqMan MicroRNA Assay (either from U6 or miR-124-3p), 1.33 µl from RT reaction, and 10 µl of TaqMan™ Fast Advanced Master Mix were mixed in a final volume of 20 µl. The PCR reaction settings were: 10 min at 95°C followed by 40 cycles of 15 sec at 95°C and 1 min at 60°C. To calculate miR-124-3p levels, the 2^-(ΔΔCt)^ method was used again, where U6 snRNA represented the housekeeping gene. GraphPad Prism 9 was used as described for RT-qPCR analysis.

### Sequencing

Prior RNA-seq, RNA quality was determined by using the Agilent RNA 6000 Nano kit (5064-1511) in an Agilent 2100 Bioanalyzer machine. Samples selected for sequencing had a RIN value > 7. RNA-seq from time course data, including smNPC, mDANs and astrocytes samples, was done using the TruSeq Stranded mRNA library prep kit, single-end 75 bp read length, and a NextSeq500 machine.

RNA-seq from KD samples was done using an Illumina stranded mRNA library prep. Kit, paired-end 50 bp read length, and a NovaSeq6000 machine.

ATAC-seq samples were sequenced on a NextSeq500 machine using paired-end 75 bp read length.

ChIP-seq samples were sequenced on a NextSeq500 machine using single-end 75 bp read length.

### RNA-seq analysis

For the time course RNA-seq, including smNPC, mDANs at days 15, 30 and 50 of differentiation, non-mDAN at days 15 and 50 of differentiation, and astrocytes at day 65 of differentiation the following tools were used. Raw fastq files were assessed for quality using FastQC (https://www.bioinformatics.babraham.ac.uk/projects/fastqc/). Summary of sample quality controls was obtained using MultiQC (30). Next, the Paleomix pipeline was used for trimming sequencing adapaters using AdapterRemoval (31, 32). After adapter removal, ribosomal RNA was filtered from the data using SortMeRNA (33). Alignment to the reference genome was done using STAR (34). BAM files were validated using Picard (https://broadinstitute.github.io/picard/). Quality reads (≥Q30) were filtered using SAMtools (35). Gene counts were obtained using FeatureCounts from the Rsubread package (36). Differential expression analysis was done using the R package DESeq2 (37). For more information about the specific versions and settings used for the different tools, please refer to our repository (RNA-seq folder, RNA-seq_DataAnalysis_TimeCourse.rmd script). Genome version and annotation were GRCh38 patch 12 and Gencode human release 31, respectively.

For the RNA-seq data from the different TF knock-down experiments, including LBX1 KD, NHLH1 KD and NR2F1/2 KD, a snakemake pipeline was used (38). This pipeline includes the tools STAR, SAMtools, FastQC, FastQ Screen, AdapterRemoval, Rsubread, DESeq2, ggplot2 and apeglm (39–41). For more details about the pipeline, please refer to our repository (RNA-seq folder, RNA-seq_DataAnalysis_TF_KDs.rmd script). Genome version and annotation were GRCh38 release 102.

### ATAC-seq analysis

For the time course ATAC-seq, including smNPC, mDANs on days 15, 30, and 50 of differentiation, non-mDAN on days 15 and 50 of differentiation, and astrocytes on day 65 of differentiation the following tools were used. Raw fastq files were assessed for quality using FastQC. A summary of sample quality control was obtained using MultiQC. Using the Paleomix pipeline, trimming of sequencing adapters was done using AdapterRemoval and mapping to the reference genome with BWA (42). BAM files were validated using Picard. Quality reads (≥Q30) were filtered using SAMtools. Peak calling was performed using Genrich (https://github.com/jsh58/Genrich). For more information about the specific versions and settings used for the different tools, please refer to our repository (README.md file inside the ATAC-seq folder). Genome version was GRCh38 patch 1.

### ChIP-seq analysis

For H3K27ac ChIP-seq, including smNPC, mDANs at days 30 and 50, and non-mDANs at day 50, the following tools were used. After merging R1 and R2 raw fastq files as described by Active Motif, the new fastq files were assessed for quality using FastQC. Using the Paleomix pipeline, trimming of sequencing adapters was done using AdapterRemoval and mapping to the reference genome with BWA. BAM files were validated using Picard. Quality reads (≥Q30) were filtered using SAMtools. New BAM files were sorted according to mapping position using SAMtools prior the molecular identifier de-duping step. For de-duping, a perl script provided by Active Motif was used (not provided, it should be requested from the manufacturer. Script name rmDupByMids.pl.txt, version from 2019). For calling enhancers and super-enhancers HOMER was used (43). For more information about the specific versions and settings used for the different tools, please refer to our repository (ChIP-seq folder, Lowinput_ChIP-seq_analysis.rmd script). Genome version was GRCh38 patch 12.

### EPIC-DREM analysis

EPIC-DREM was applied as a snakemake pipeline. As an input, pre-processed BAM files from the ATAC-seq analysis and the gene counts from the previously described RNA-seq analysis were used, including smNPC, and mDANs at days 15, 30 and 50. ATAC-seq peak calling was performed with Genrich over replicates. The Regulatory Genomics Toolbox was used to identify footprints in called peaks (21) and subsequently, TF-gene affinities were calculated using TEPIC (22). The resulting time-point-specific lists of TF-gene links were merged and filtered according to expression, removing links from unexpressed TF. TF was considered unexpressed if it presented with a transcripts per million (TPM) value < 1 in all analyzed time points. Time-point-specific GRN were identified with interactive Dynamics Regulatory Events Miner (iDREM) (44). Results from iDREM were further processed in R for visualization and GO enrichment was performed using clusterProfiler (45). For more details about the different parameters used, please refer to our repository (EPIC-DREM folder).

### Primers

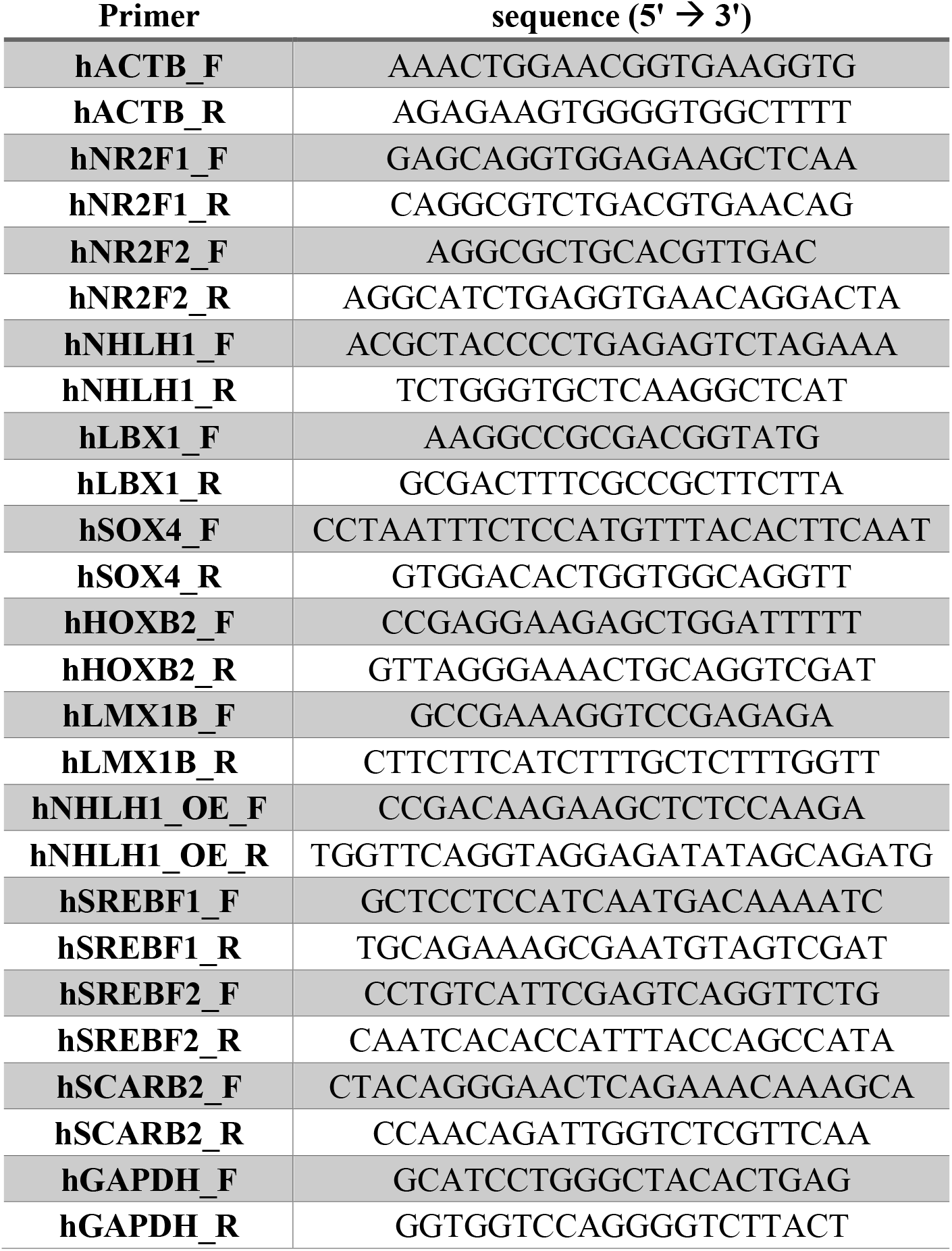

### Antibodies

| Antigen | Specie | Dilution | Cat No |
| --- | --- | --- | --- |
| TH | Rabbit | 1:250 | Sigma T8700-1VL |
| NeuN Alexa fluor 488 conjugated | Mouse | 1:250 | Sigma MAB377X |
| Histone H3 acetyl K27 (H3K27ac) | Rabbit | ** | Abcam Ab4729 |
| Rabbit IgG Alexa fluor 647 | Goat | 1:1000 | Invitrogen A27040 |

## Bacterial glycerol stocks

All bacterial glycerol stocks are from GeneCopoeia. The different shRNA constructs used were:

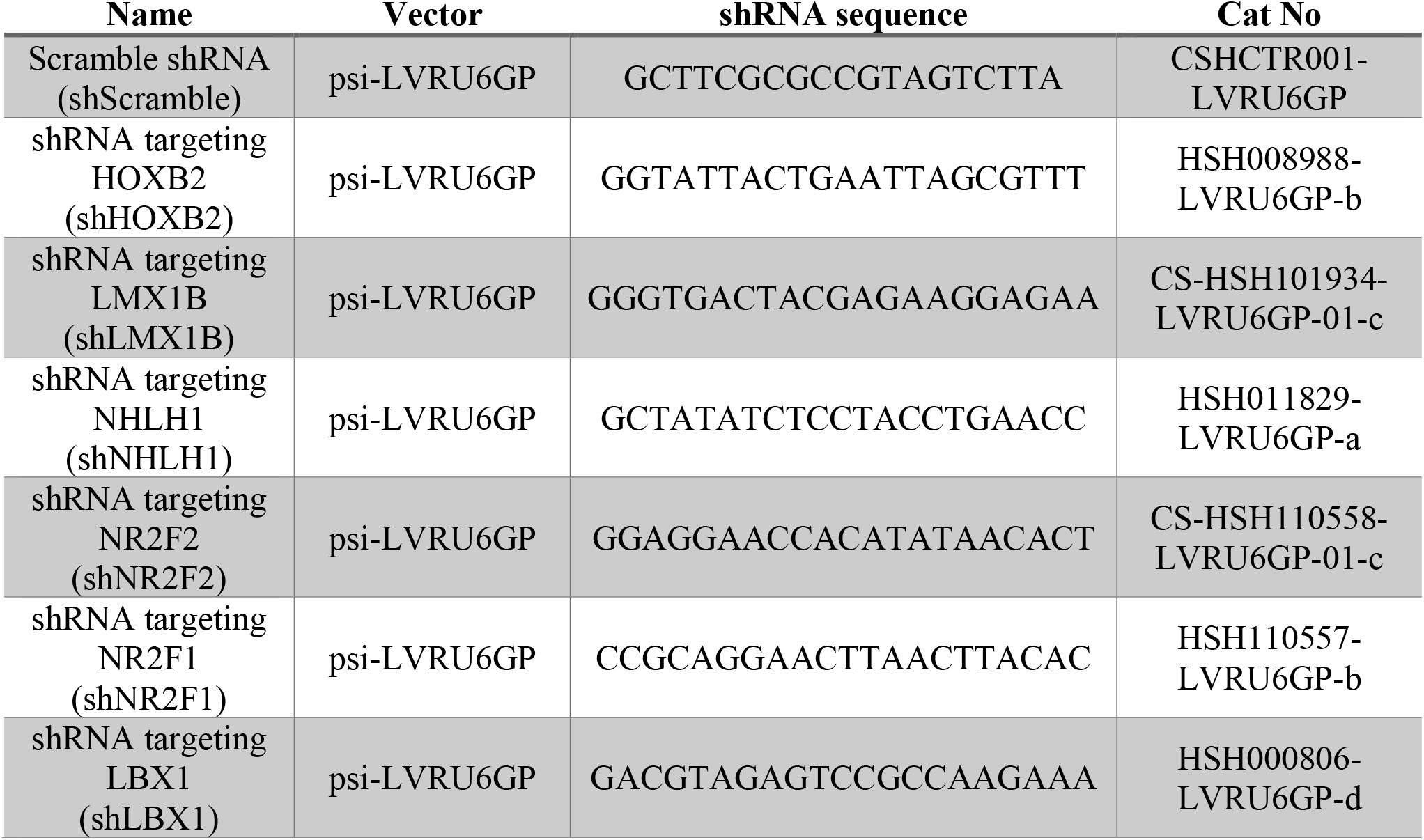

The different overexpression constructs used were:

| Name | Vector | Accession No | Cat No |
| --- | --- | --- | --- |
| Control empty vector (GFP OE) | pEZ-Lv228 | ** | EX-NEG-Lv228 |
| NHLH1 overexpression (NHLH1 OE) | pEZ-Lv228 | NM_005598 | EX-F0569-Lv228 |
| LBX1 overexpression (LBX1 OE) | pEZ-Lv228 | NM_006562 | EX-T1288-Lv228 |
| NR2F2 overexpression (NR2F2 OE) | pEZ-Lv228 | NM_021005 | EX-C0221-Lv228 |

## Results

### Generation of paired transcriptomic and chromatin accessibility profiles of mDAN differentiation

mDANs were enriched for transcriptomic and epigenomic profiling based on the expression of P2A-mCherry reporter, stably expressed under the control of the endogenous promoter of the TH gene, coding for tyrosine hydroxylase, the rate-limiting enzyme for dopamine biosynthesis (Figure 1A, (19)). The presence of the mCherry reporter allowed the purification of mDANs from heterogeneous iPSC-derived neuronal cultures by FACS at multiple time points of differentiation (Figure 1A-B). Reporter expression correlated with TH expression, as validated by immunocytochemistry (Figure 1B). Using the reporter cell line, paired transcriptomic and chromatin accessibility profiles were generated from neuronal progenitor cells (smNPCs) and mDANs after 15, 30, and 50 days of differentiation, as shown in Figure 1A. Time points were selected based on culture features reported by (46): mDANs appear in the culture by day 15, after the cells are incubated in maturation medium (Figure 1A); they start to show electrophysiological activity after 30 days of differentiation (46). Finally, by day 50 mDANs begin to resemble more mature neurons. Gene expression and chromatin accessibility profiles were generated from iPSC-derived astrocytes differentiated from the same reporter cell line for 65 days (Figure 1, (26)) as additional reference data.

The data generated from mDANs showed cell-type-specific gene expression and chromatin accessibility profiles based on established markers of these cells (TH, EN1, DDC, and LMX1B) when compared to non-mDANs, and in contrast to either pluripotency/early neuroectoderm markers (PAX6) or glial markers (GFAP) (Figure 1C, (14)(47)(48)). The transcriptome analysis of the mCherry population across differentiation confirmed a downregulation of pluripotency genes in parallel with induction of pan-neuronal markers and, importantly, mDAN-specific marker genes (Supplementary Figure 1, (13, 49)). Moreover, some of the genes selective for either A9 and A10 mDAN subtypes became upregulated, while others remained undetected, suggesting that the cells have not adopted a subtype-specific identity. Principal component analysis (PCA) confirmed a clear separation of smNPCs, mDANs and astrocytes at the level of both transcriptome and chromatin accessibility (Figure 1D). Taken together, our reporter cell line and the obtained genome-wide profiles can be used for further characterization of the regulatory landscape of mDAN differentiation.

### Integrative analysis predicts key regulators of human mDAN differentiation

To integrate the paired transcriptomic and epigenomic profiles to identify key regulators of mDAN differentiation, our previously published EPIC-DREM pipeline was adapted for use with ATAC-seq data and applied (Figure 1E, (8)). In short, genome-wide accessible regions were determined per cell type and time point, and footprinting analysis was performed to predict TF binding sites (TFBS) in these regions using HINT-ATAC (21). Computed TF-gene scores based on accessibility signal and TF binding strength were integrated to finally define regulators across time in previously described statistical framework (22, 50). The time point-specific predictions were combined with the time series gene expression changes by DREM (23) to build a time point-specific gene regulatory network of mDAN differentiation. DREM identifies gene expression bifurcation points across the time points analyzed, corresponding to groups of co-expressed genes. For each bifurcation point (also called split node), TFs are ranked according to the highest number of target genes within the group based on accessibility data predictions. Therefore, DREM highlights the TFs controlling most of the observed transcriptional changes across time.

Figure 2A shows the result of EPIC-DREM on mDAN time course data. There are a total of 26 split nodes with a total of 327 TFs ranked differently across them (Supplementary Table 1). Highlighted in red are different top-ranked TFs known to be involved in the regionalization, differentiation, and specification of mDANs. For example, well-described TFs controlling mDAN differentiation, such as NR4A2 (also known as NURR1), LMX1A/B and EN1, were identified among the main regulators (51– 54). Pioneer TFs, critical for cell reprogramming due to their ability to alter chromatin structure (55), such as ASCL1 and NEUROG2, also appeared as top-ranked TFs (56, 57). Other factors more involved in the regionalization and differentiation of the midbrain such as GBX2, NKX6-1, PBX1, SREBF1, NR1H2 (also known as LXRβ), and MSX1 were captured by EPIC-DREM (58–62). Thus, our EPIC-DREM predictions are in line with the existing literature.

**Figure 2:**
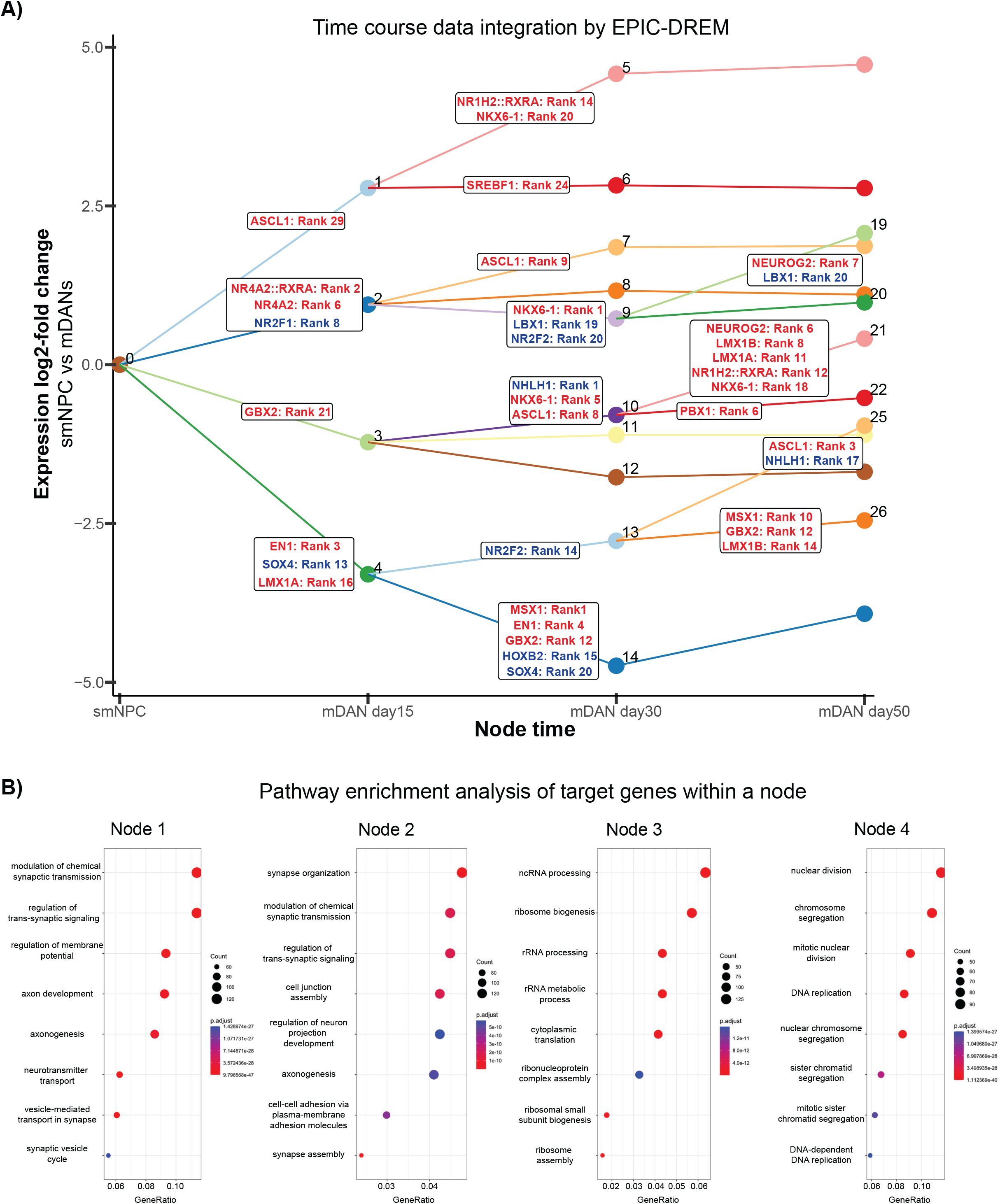
Integrative analysis of time series transcriptomic and chromatin accessibility data predicts key regulators of mDAN differentiation. **A)** EPIC-DREM results from the integration of time course data on gene expression and chromatin accessibility profiles of mDAN differentiation from Figure 1. X-axis represents the time points analyzed and y-axis represents expression log_2_-fold changes across time. EPIC-DREM result contained a total of 26 split nodes with an associated list of TFs ranked according to their regulatory importance for the genes contained within the node. See also Supplementary Table 1. Highlighted in red are TFs ranked as top regulators and previously associated to control of mDAN differentiation. Highlighted in blue are the novel identified TFs that were selected for functional analysis. **B)** Pathway enrichment analysis for the genes contained in the first 4 nodes created by EPIC-DREM captures the main biological processes regulated at early stages of neuronal differentiation.

Moreover, pathway enrichment analysis of the genes included in the first four split nodes revealed enrichment for neuronal functions such as axon development and synapsis formation among the upregulated genes (Figure 2B, nodes 1 and 2) and enrichment for ribosomal RNA production and cell division for downregulated genes (Figure 2B, nodes 3 and 4, respectively), consistent with switching from the multipotent state to neuronal differentiation. Thus, EPIC-DREM can highlight the main biological processes governing mDAN differentiation.

### Identification of key TFs controlled by super-enhancers

While EPIC-DREM allowed us to predict the TFs controlling the highest number of target genes during mDAN differentiation, we set out to further prioritize these TFs by identifying those that are themselves under high regulatory load in a cell type-selective manner (4, 63). To do this we performed low input ChIP-seq for H3K27ac, a histone mark associated with active enhancers, for mDANs at days 30 and 50 of differentiation to determine which TFs display association with SEs at the respective time point. For the association, SE regions were overlapped with the genomic coordinates of the genes encoding for TFs and expressed in a time point-specific manner to obtain a list of 49 TFs controlled by SEs across both time points (Figure 3A, Supplementary Table 2). Finally, the list of all SE-associated TFs at either time point was compared with a list of TFs combining the top 20 ranked TFs from each of the split nodes from EPIC-DREM (Figure 3B, Supplementary Table 2). Among the 49 SE-associated TFs, 17 were also among the top 20 ranked TFs from EPIC-DREM (Figure 3B, Supplementary Table 2).

**Figure 3:**
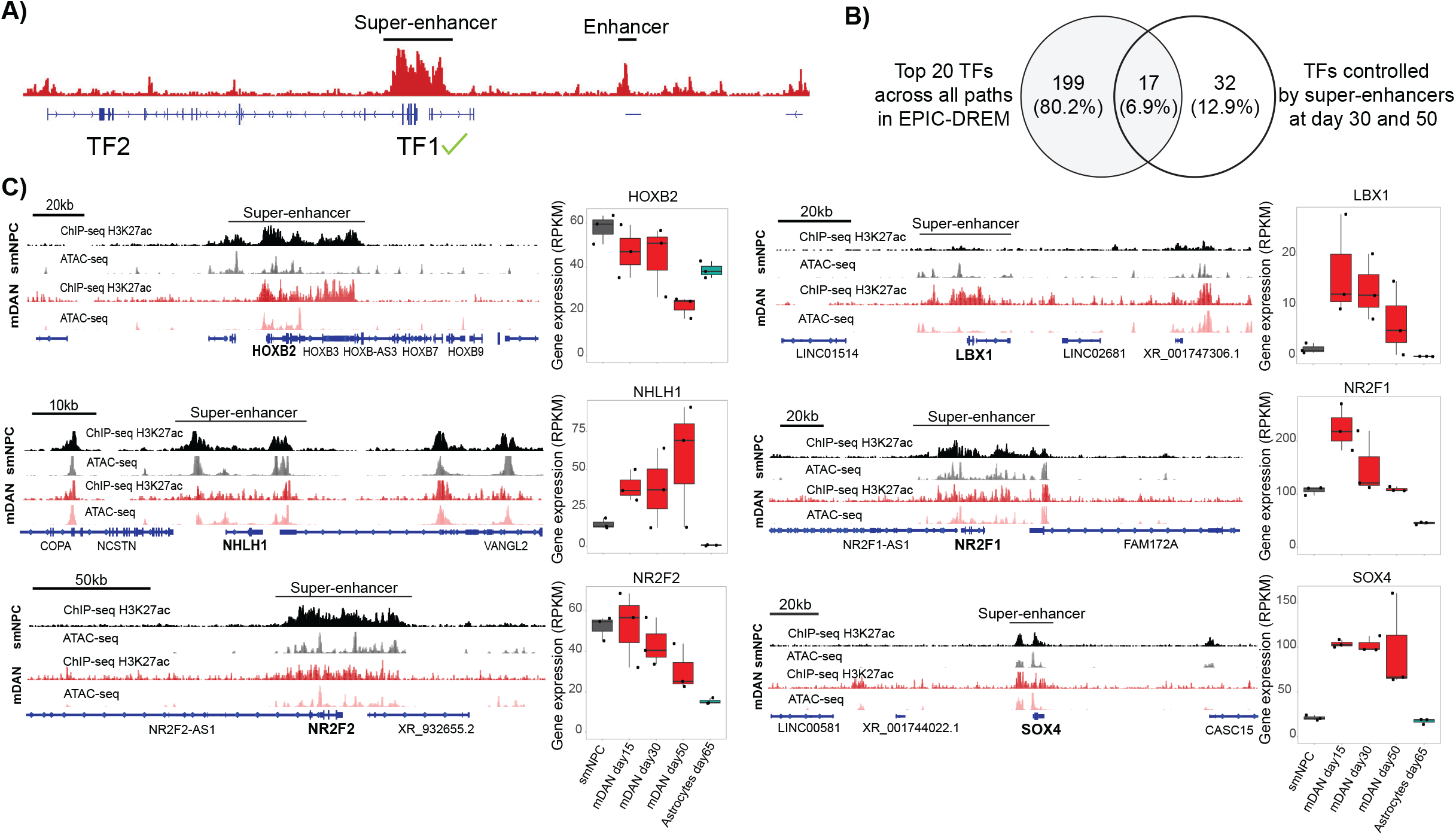
Identification of key TFs controlled by super-enhancers. **A)** Schematic representation of the selection of TFs controlled by super-enhancers. First, TF has to be expressed at the specific time point of analysis and second, its gene body had to be located under a super-enhancer region in order to be included. **B)** Venn analysis of the top 20 TFs across all split nodes from EPIC-DREM with the list of TFs controlled by super-enhancers across the analyzed time points. **C)** H3K27ac signal and chromatin accessibility profiles at the loci of the novel candidate TFs, highlighting the identified super-enhancer regions in mDANs, together with the expression dynamics during mDAN differentiation. ATAC-seq and ChIP-seq tracks are plotted under the same scale per dataset for comparison purposes.

The list of 17 TFs was further explored to select the most promising candidates for functional analysis. For this, a literature search was performed to see whether these TFs had already been associated with mDAN function or development. For example, as TCF4 and MEIS1 have been previously related to mDAN subset specification and striatal dopaminergic system formation, respectively, and consequently were not included in follow-up experiments (64, 65). From the remaining TFs, HOXB2, LBX1, NHLH1, NR2F1 (also known as COUP-TFI), NR2F2 (also known as COUP-TFII) and SOX4 were found to present the strongest SE signals and most dynamic gene expression profiles and, therefore, were selected for functional analysis as novel candidate regulators of mDAN differentiation (Figure 3C).

Among the candidates, HOXB2 has been mainly associated with neural crest and hindbrain patterning (66). No relation of HOXB2 with mDANs has been described before. HOXB2 is clustered with other HOXB genes that are also proximal to the same SE. Most of the chromatin accessibility is observed within the HOXB2 gene body (Figure 3C). HOXB2 was abundantly expressed in smNPCs and early time points of differentiation but decreased in day 50 mDANs, in keeping with its predicted role as a regulator of downregulated genes (Figure 2A, Supplementary Table 1).

LBX1 has been implicated in the development of different types of neurons such as retrotrapezoid nucleus (RTN) neurons, GABAergic neurons, and somatosensory interneurons in the spinal cord (67– 69), highlighting a role for this TF in cell fate decisions. LBX1 was the only TF among the selected candidates to present an mDAN-specific SE (Figure 3C). Consistently, LBX1 expression was induced >10-fold upon early mDAN differentiation.

NHLH1 is mainly characterized as an early pan-neuronal marker that determines many neuronal cell fate decisions (70–73). NHLH1 expression was continuously increased during differentiation with almost 70 RPKM detected in day 50 mDANs (Figure 3C).

Temporal expression dynamics of NR2F1 and NR2F2 in the developing brain are important for the specification and balance of different neuronal lineages as well as neuron-to-glia transitions (74–77). Recently, NR2F1 was found to be deregulated in iPSCs from PD patients carrying the LRRK2-G2019S mutation (29). During early mDAN differentiation NR2F1 expression increased to particularly high levels (>200 RPKM), while NR2F2 expression decreased in later time points compared to smNPCs (Figure 3C).

SOX4, like many SOX genes, has been described to be implicated in neurogenesis and maintenance of progenitor cells during development. Interestingly, the role of this TF in generating TH-expressing cells from sympathetic or enteric nervous systems has been described (78, 79). Also, SOX4 expression increased to >100 RPKM during mDAN differentiation and was accompanied by an increased enhancer signal at the locus (Figure 3C).

Overall, these candidates presented great potential as cell fate and differentiation determinants with no previous relation to mDANs.

### LBX1, NHLH1, and NR2F1/2 are necessary for mDAN neurogenesis

To functionally validate the data-driven predictions *in vitro*, TFs were knocked down (KD) during differentiation to determine their impact on the mDAN number in mixed cultures. A lentiviral vector containing a shRNA targeting the different candidate TFs and a GFP reporter was used to control for transduction efficiency. To assess the effect of the TFs, two different experimental designs for transduction were used: early and late transduction (Figure 4A). For early transduction, cells were transduced on day one of differentiation while for late transduction, cells were transduced on day nine, just before they entered the maturation phase. In both cases, the cells were analyzed on day 15 (Figure 4B shows the results from the KD experiments). LMX1B served as a positive control as its role in mDAN differentiation is well characterized (53, 80). SOX4 was omitted as we were unable to identify an shRNA with a KD efficiency of >25%. As expected, LMX1B KD reduced mDAN numbers in the cultures as assessed by the mCherry reporter signal (Figure 4B). Upon early transduction, LMX1B KD reduced the numbers of mDAN in the culture by ∼60%. However, the KD efficiency for LMX1B, as measured by RT-qPCR, was very variable, which can be explained by the low levels of GFP positive cells (∼40%) upon analysis on day 15 of differentiation (Supplementary Figure 2A), possibly masking the KD of this TF by the GFP negative cells in the culture. On the other hand, with late transduction, good transduction efficiency and strong KDs were observed on the day of analysis for LMX1B. Interestingly, late KD of this TF reduced the mDAN numbers by only 30% but also decreased the overall cell density in the cultures. This highlights the dual role of LMX1B during differentiation and correlates with previous studies (80), emphasizing the biological relevance of our cellular system.

**Figure 4.**
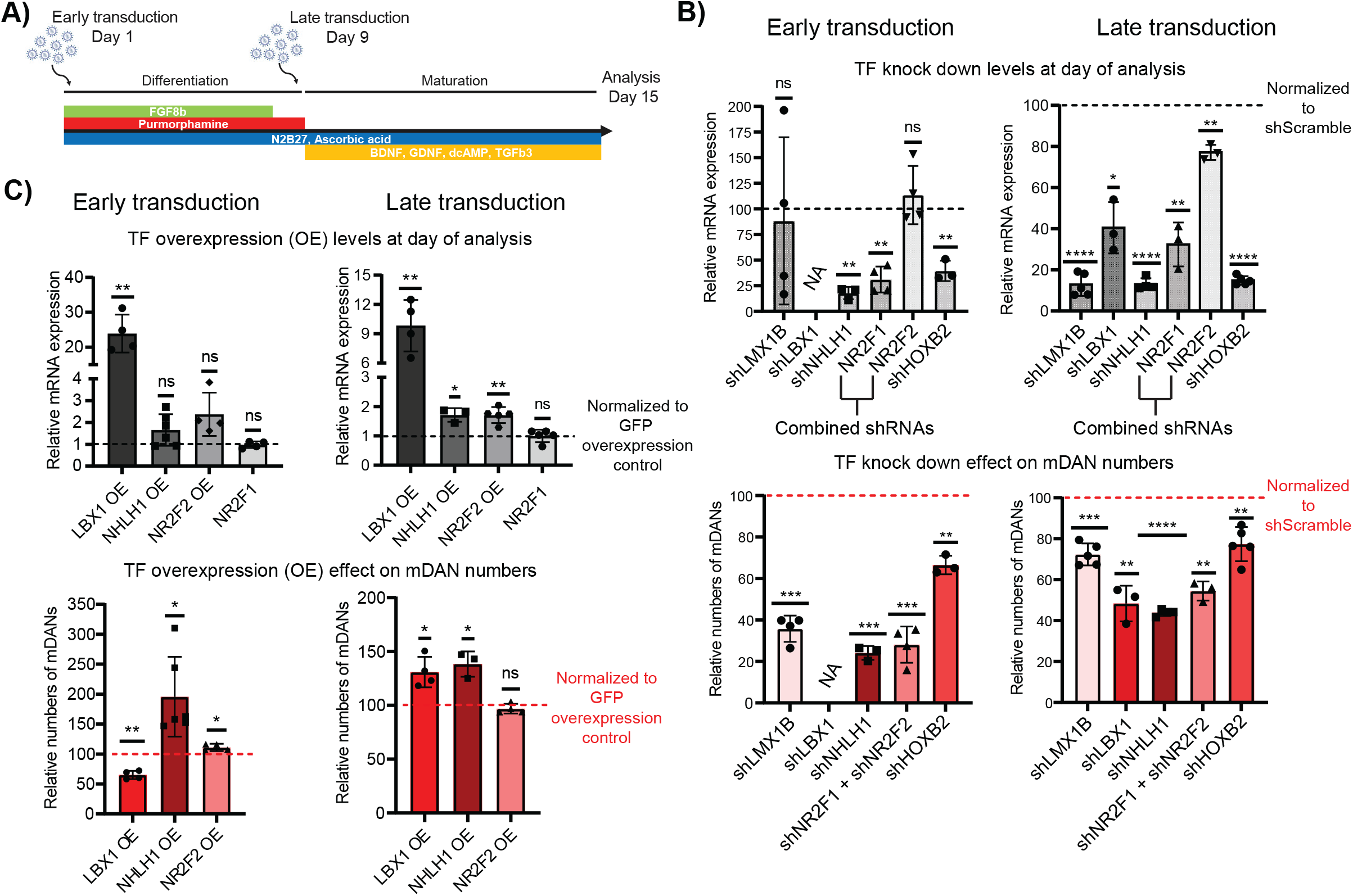
LBX1, NHLH1, and NR2F1/2 are necessary for mDAN differentiation and LBX1 and NHLH1 can also improve mDAN neurogenesis. **A)** Graphic representation of the two different transduction approaches (day 1 and day 9) used in this study and the day of analysis (day 15) for all samples. **B)** TF knock down results for early and late transduction of shRNA lentiviral particles. mRNA levels were always normalized to shScramble per replicate. Relative mDAN numbers were calculated based on mCherry signal and normalized to the mCherry population of the shScramble condition per replicate. One sample t-test was used for statistical analysis, taking 100 as the theoretical mean. **C)** TF overexpression results for early and late transduction of lentiviral particles containing codon optimized cDNA. Expression was normalized to GFP overexpression per replicate. Relative mDAN numbers were also calculated according to mCherry signal but normalized to GFP overexpression per replicate. One sample t-test was used for statistical analysis, taking 1 and 100 as the theoretical mean for gene expression and for mDAN numbers, respectively. For knock-down and overexpression experiments, N ≥ 3. * = p-value < 0.05, ** = p-value < 0.01, *** = p-value < 0.001, **** = p-value < 0.0001, and ns = not significant.

LBX1 KD by early transduction was found to severely decrease the cell numbers, with very few cells remaining by day 5-6 post transduction, preventing a more detailed analysis of the effect of this TF on mDANs (Supplementary Figure 2A). Although LBX1 KD by late transduction also resulted in some decrease in cell numbers, the surviving population displayed good transduction efficiencies and allowed us to study the impact of LBX1 on mDANs. After late transduction, LBX1 KD efficiency on the day of analysis was ∼60% and mDAN numbers were reduced by 50% (Figure 4B).

NHLH1 KD had the strongest effect on mDAN numbers among the candidates tested. KD efficiencies in both early and late transduction were around 80% and mDAN numbers were reduced by roughly 80% and 60% in early and late transduction, respectively. Although the total number of cells was not affected by the NHLH1 KD in early transduction, GFP-positive cells represented only half of the cells on the day of analysis, similar to what was observed for LMX1B KD (Supplementary Figure 2A).

Since NR2F1 and NR2F2 are known to act redundantly (74), lentiviral particles expressing the respective shRNAs were combined to target both TFs to avoid possible compensatory mechanisms. While individual KD of NR2F1 or NR2F2 was successful (data not shown), in dual KD conditions only NR2F1 levels were significantly reduced (Figure 4B). Nevertheless, the combined NR2F1/2 KDs markedly affected mDAN differentiation, reducing mDAN numbers in the culture by close to 80% and 50% in early and late transduction, respectively. In addition, when NR2F1/2 were KD, cells could keep good transduction efficiencies until the day of analysis despite the negative effect on mDAN numbers, in contrast to what was observed for LMX1B and NHLH1 KDs (Supplementary Figure 2A).

Lastly, for HOXB2 good transduction efficiencies were observed on the day of analysis, with >60% KD efficiency in both early and late transduction. However, although HOXB2 KD reduced the numbers of mDANs in the culture, the effect was the weakest among the tested TFs.

In summary, LBX1, NHLH1 and NR2F1/2 were all found to be necessary for mDAN differentiation. Their depletion during the specification of these neurons exhibited a more robust phenotype than that of LMX1B, a well-established regulator of mDAN differentiation, thereby demonstrating an important regulatory role for LBX1, NHLH1, and NR2F1/2. On the other hand, loss of HOXB2 only had a limited effect on mDAN levels.

### Elevated expression of LBX1 or NHLH1 can increase mDAN numbers

To further determine the role of the identified TFs in mDANs, LBX1, NHLH1 and NR2F1/2 were overexpressed during differentiation using the same lentiviral transduction approach as in the earlier KD experiments. Figure 4C shows the results from the overexpression experiment for the three different TFs. For LBX1, strong overexpression (9-20-fold compared to control vector) and high levels of transduced cells were observed on the day of analysis (Supplementary Figure 2B) for both early and late transduction. However, the induction of LBX1 had opposite effects depending on the time point of differentiation. While LBX1 overexpression early during differentiation had a negative impact on mDAN numbers, late overexpression resulted in significantly increased mDAN numbers (Figure 4C).

Overexpression of NHLH1 during early differentiation resulted in an almost 2-fold increase in mDANs, despite a modest increase of gene expression levels by 50% and reduced number of transduced cells based on the GFP signal on the day of analysis. With late induction of NHLH1 expression, again, high levels of transduced cells, and stable overexpression could be observed, leading to a significant increase in mDAN numbers.

We were unable to achieve any meaningful overexpression of NR2F1, most likely due to its already very high endogenous expression levels (Figure 3C and data not shown). Therefore, only NR2F2 overexpression was tested. While the expression fold change upon NR2F2 overexpression was comparable to that achieved for NHLH1, this had almost no effect on the cultures. Early transduction showed a very minor, albeit significant increase in mDAN levels, while late transduction for NR2F2 overexpression had no observable effect on mDAN numbers. Interestingly, NR2F2 overexpression could not induce NR2F1 expression (Figure 4C).

Taken together, LBX1 and NHLH1 were able to increase the number of mDAN, with the timing of overexpression being the key to producing this effect for LBX1. NHLH1 showed again the strongest effect on mDAN numbers, as also observed for its KD, and the effect was independent of the developmental timing. Lastly, NR2F2 overexpression did not present any benefit regarding mDAN neurogenesis. This is likely due to the considerable levels of expression by the endogenous and functionally redundant NR2F1.

### NHLH1 controls miR-124-3p expression in mDANs

To further characterize the role of NHLH1 in mDAN differentiation, RNA-seq was performed using the samples from the late transduction KD experiments (Figure 4A). NHLH1 KD led to a total of 491 differentially expressed genes (DEGs) (absolute log_2_-fold change > 1, FDR < 0.05) (Figure 5A, Supplementary Table 3). Using an Ingenuity Pathway Analysis (IPA) for upstream regulators explaining the transcriptional changes produced by NHLH1 KD (Figure 5B, top 10 upstream regulators based on z-score, p-value < 0.05) predicted miR-124-3p to be downregulated and to belong to the key regulators driving the expression changes (81). Consistently, g:Profiler (82) analysis of upregulated genes from NHLH1 KD predicted miR-124-3p to control many of the genes (Supplementary Table 3). Moreover, miR-124-3p was the only regulator predicted by both methods, IPA and g:Profiler. miR-124 is the most abundant microRNA in the brain with neurogenic properties and has been associated with dopaminergic neurodegeneration in PD (83). Indeed, further exploration of our epigenomic data from mDANs confirmed the three loci coding for miR-124 (MIR124-1 to -3) to be highly accessible with large regions occupied by H3K27ac (Supplementary Figure 3A). Moreover, the primary transcripts of miR-124-1 and miR-124-2 presented an increased expression specifically in mDANs (Supplementary Figure 3A). To validate the predicted reduction in miR-124 expression upon NHLH1 depletion, a TaqMan microRNA assay was used. Importantly, a strong and significant downregulation of mature miR-124-3p could be confirmed (Figure 5C).

**Figure 5:**
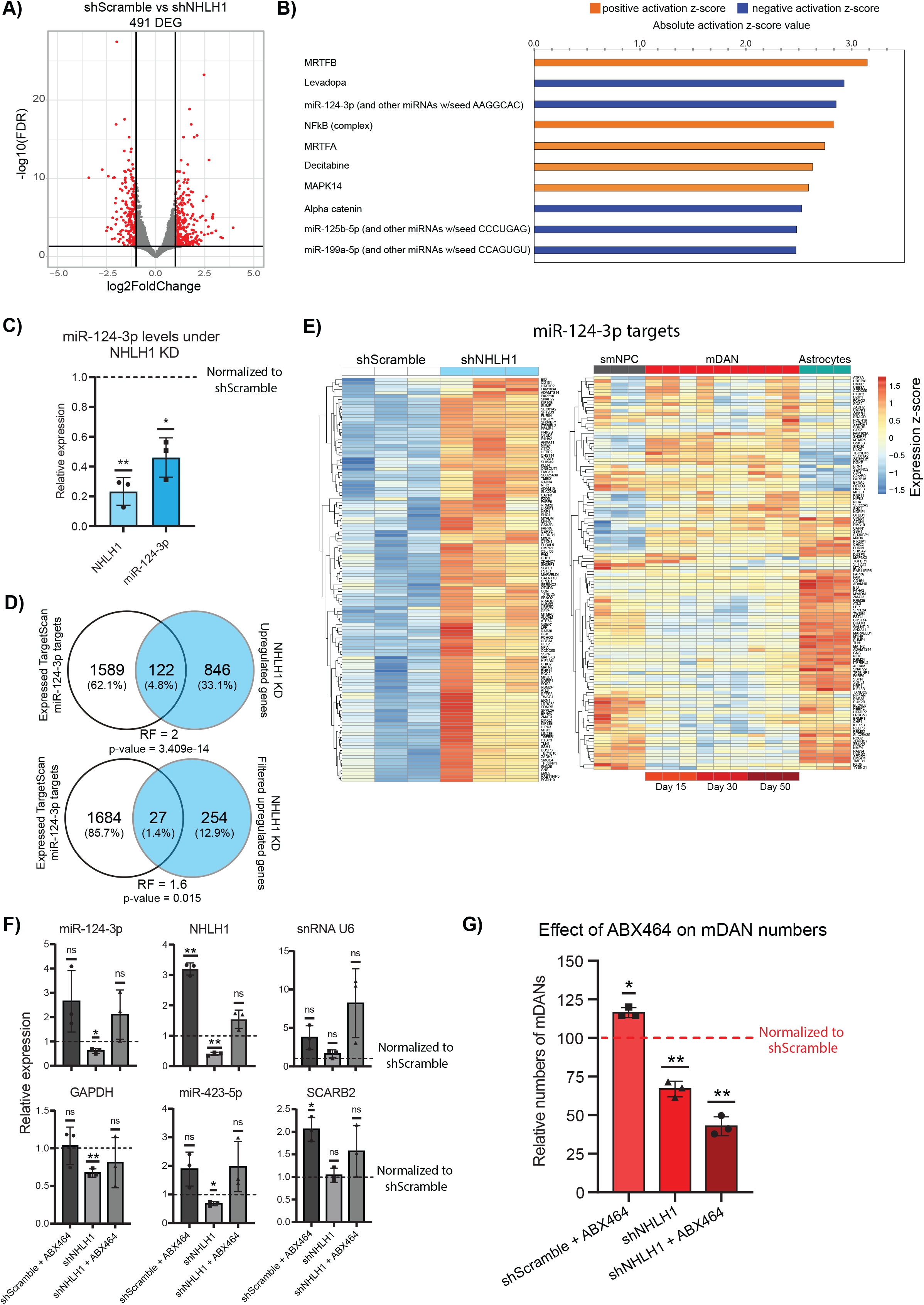
NHLH1 controls mDAN differentiation by regulating miR-124-3p expression. **A)** Volcano plot showing DEGs from RNA-seq analysis of neurons at day 15 following a late transduction with shScramble or shNHLH1. N = 3. Black lines represent cut-off according to FDR < 0.05 and absolute log_2_-fold change > 1. **B)** Ingenuity Pathway Analysis (IPA) predicted top 10 upstream regulators based on absolute Z-score value using DEGs from panel A. P-value < 0.05 for all regulators. **C)** Bar plots showing results from TaqMan assay determining miR-124-3p levels upon NHLH1 KD. Expression was normalized to shScramble samples. One sample t-test was used for statistical analysis, taking 1 as the theoretical mean. N = 3 **D)** Venn analysis of predicted and expressed miR-124-3p targets from TargetScan with upregulated genes upon NHLH1 KD. Two overlaps were done, one using the upregulated genes with a p-value < 0.05 and the second with upregulated genes with a log_2_-fold change > 1 and p-value < 0.05. For overlaps, a hypergeometric test was used to determine statistical significance. RF = representation factor. RF > 1 indicates more overlap than expected by chance. **E)** Heatmaps showing the expression of the 122 predicted miR-124-3p targets upregulated upon NHLH1 KD. Genes were plotted in the RNA-seq data from the KD experiment and the time course data containing smNPC, mDANs and astrocytes. **F)** RT-qPCR and TaqMan assays were used to determine the expression of different genes and microRNAs upon ABX464 treatment and NHLH1 KD conditions. Expression values were all normalized to ACTB. Normalization between groups was done using shScramble as the reference condition. One sample t-test was used for statistical analysis, taking 1 as the theoretical mean. N = 3. **G)** Effect of ABX464 treatment and NHLH1 KD conditions on mDAN numbers based on mCherry signal. mDAN numbers were normalized to shScramble. One sample t-test was used for statistical analysis, taking 100 as the theoretical mean. N = 3. * = p-value < 0.05, ** = p-value < 0.01, and ns = not significant.

Taking advantage of TargetScan (https://www.targetscan.org), a database containing predicted microRNA targets (84), a list of predicted targets for miR-124-3p was filtered for the genes also expressed in our RNA-seq data (Supplementary Table 3). The obtained gene list was compared with the list of upregulated DEGs from NHLH1 KD to determine how many of the genes could be affected by the downregulation of miR-124 (Figure 5D). Strikingly, over 12% of all upregulated genes belonged to primary targets of miR-124, a significantly larger proportion than expected by chance (hypergeometric test, p-value = 3.409e-14). Plotting the 122 targets of miR-124-3p in the KD RNA-seq data confirmed a strong upregulation, as expected (Figure 5E). When the expression of the same targets in smNPCs, across mDAN time course, and astrocytes was plotted, it became clear that most of the microRNA targets are enriched in astrocytes or smNPCs (Figure 5E). Overall, the results highlight miR-124 as a likely mediator of NHLH1-controlled gene regulation, contributing to mDAN specification.

To directly test the contribution of miR-124, a small molecule treatment was used to stimulate miR-124 expression during differentiation and determine whether there would be any benefit for mDAN differentiation. Recent studies have described a new role for ABX464, a quinoline with antiviral properties, in the selective and specific induction of miR-124 (85–87). This molecule binds to the Cap binding complex (CBC) at the 5’ end of the primary transcript and promotes the selective splicing of LINC00599, the host gene of miR-124-1. We found that ABX464 could induce miR-124-3p by around two-fold, but the required concentration had a negative effect on cell density in culture (Supplementary Figure 3B). Nevertheless, as small molecules can be powerful tools for their use in biomedicine, and their optimization is possible by chemical modifications, we proceed with the testing ABX464 during mDAN differentiation. ABX464 was added to the cells when they entered the maturation medium on day 9 of differentiation. The molecule was tested under normal differentiation and in cells transduced by shNHLH1.

Interestingly, ABX464 treatment strongly affected both mRNA and microRNA expression (Figure 5F, for original Ct values, please see Supplementary Figure 3D). Moreover, ABX464 also increased U6 snRNA, preventing its use for normalization, while another endogenous microRNA (miR-423-5p) and SCARB2 mRNA were also affected. Hence, we normalized all mRNAs and microRNAs to ACTB expression. Although not significant, ABX464 treatment appeared to increase miR-124-3p levels, while it was downregulated upon NHLH1 KD, as expected. NHLH1 expression was also upregulated upon ABX464 treatment that even reversed NHLH1 repression in NHLH1 KD cells. The rescue of NHLH1 and the increase of miR-124-3p in the cells treated with both ABX464 and shNHLH1 was not enough to rescue the mDAN loss (Figure 5G). However, ABX464 treatment alone significantly increased mDAN numbers in the culture, possibly due to increased NHLH1 and miR-124-3p expression (Figure 5F-G and Supplementary Figure 3D).

### LBX1 regulates cholesterol metabolism

LBX1 KD resulted in a total of 2241 DEGs as assessed by RNA-seq (Figure 6A, Supplementary Table 4). Next, pathway enrichment analysis of the identified DEGs was performed using IPA to determine the main processes altered by the LBX1 KD (81). Figure 6B shows the top 10 pathways affected by the KD based on the significance of the enrichment. Cholesterol biosynthesis appeared as one of the main pathways affected by LBX1 and was predicted to be reduced upon LBX1 depletion. Indeed, most genes involved in cholesterol biosynthesis according to the human metabolic reconstruction (RECON) (88), were significantly downregulated upon the KD (Figure 6C). TFs involved in cholesterol metabolism have been previously associated with mDAN development and neurogenesis (60, 62). Consistently, we found the upstream transcriptional regulators of cholesterol metabolism, SREBF1 and SREBF2 (89), to be downregulated by LBX1 KD), although only SREBF1 was reduced more than 2-fold (Figure 6D). The decreased levels were also confirmed by RT-qPCR (Figure 6E). Moreover, upstream regulators of SREBF1, NR1H3 and NR1H2 (LXRα/β) were also found to be deregulated (LXRα/NR1H3 log_2_-fold change = −0.515 and LXRβ/NR1H2 log_2_-fold change = 0.87). This is consistent with the observed pathway enrichment for LXR activation (Figure 6B). Finally, SREBF1 and NR1H2 appeared as top regulators in our EPIC-DREM analysis (Figure 2A).

**Figure 6:**
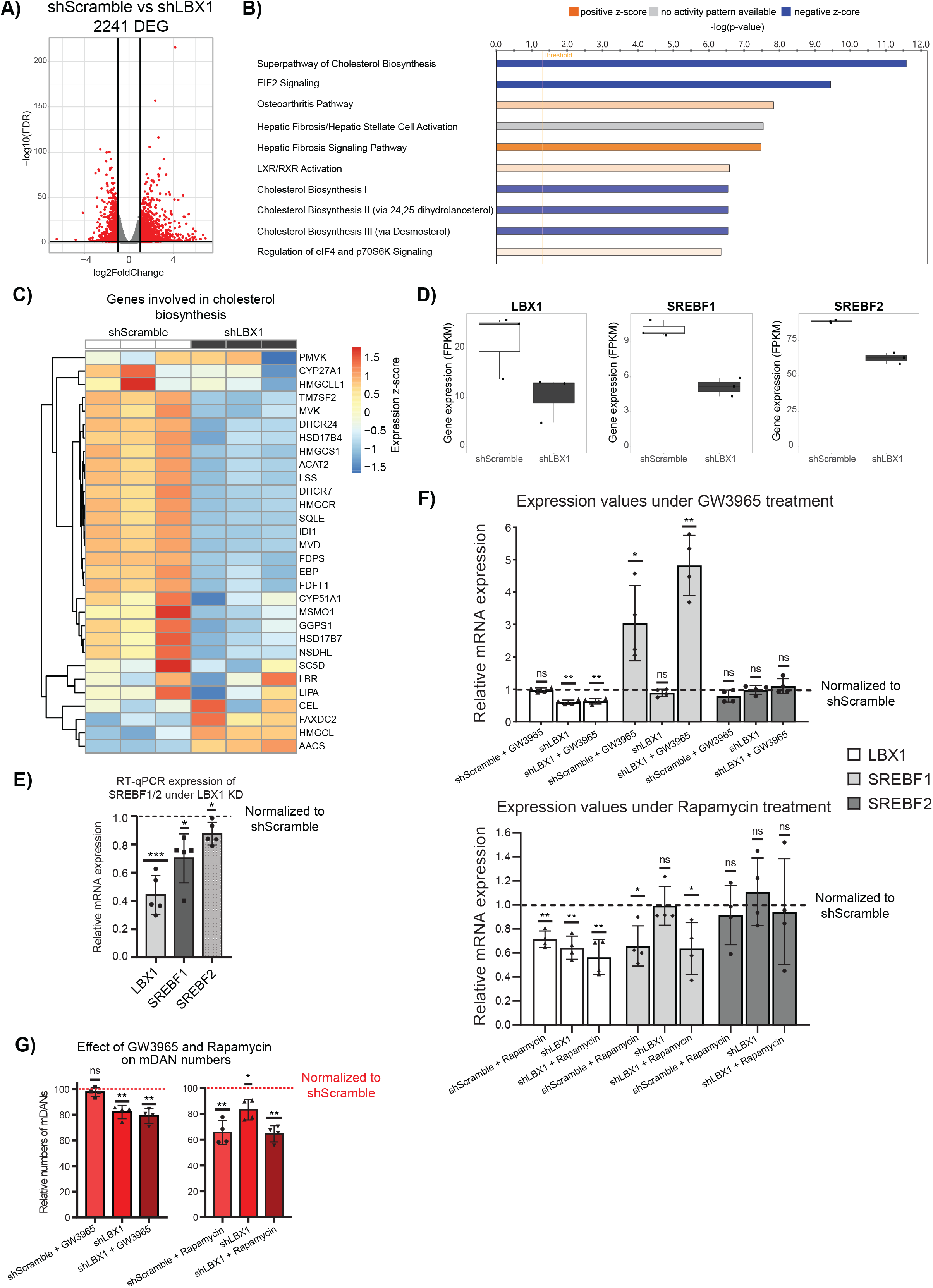
LBX1 controls cholesterol metabolism. **A)** Volcano plot showing DEGs from RNA-seq analysis of neurons at day 15 following a late transduction with shScramble or shLBX1. N = 3. Black lines represent cut-off according to FDR < 0.05 and absolute log_2_-fold change > 1. **B)** Top 10 pathways from Ingenuity Pathway Analysis (IPA) based on significance of enrichment using DEGs from panel A. X-axis represents the -log_10_ p-value of the enrichment. **C)** Heatmap showing the differences in expression between shScramble and shLBX1 of the genes involved in cholesterol biosynthesis. **D)** Expression levels (FPKM) of LBX1, SREBF1 and SREBF2 in the RNA-seq analysis. **E)** RT-qPCR validation of SREBF1/2 downregulation due to LBX1 KD in late transduced neurons at day 15 of differentiation. N = 5. mRNA levels were normalized to shScramble per replicate. One sample t-test was used for statistical analysis, taking 1 as the theoretical mean. **F)** Impact of GW3965 and rapamycin treatments on gene expression in differentiating mDANs and upon LBX1 KD. Expression of LBX1, SREBF1 and SREBF2 was normalized to shScramble per replicate. N = 4. One sample t-test was used for statistical analysis, taking 1 as the theoretical mean **G)** Effect of GW3965 and rapamycin treatments on mDAN numbers based on mCherry signal and normalized to the mCherry population of the shScramble per replicate. N = 4. One sample t-test was used for statistical analysis, taking 100 as the theoretical mean. * = p-value < 0.05, ** = p-value < 0.01, *** = p-value < 0.001, and ns = not significant.

These results suggested a novel role of LBX1 in controlling mDAN differentiation via cholesterol metabolism. Therefore, stimulating the cholesterol metabolism pathway should promote mDAN differentiation and could rescue the LBX1 KD effect. GW3965 is a potent and well-described synthetic agonist of NR1H3/2 (LXRα/β). LXR activation can lead to SREBF1 induction and improved mDAN differentiation (60). Hence, neurons were treated with GW3965 under normal differentiation or upon LBX1 KD, starting from day 9 of differentiation, and analyzed on day 15 of differentiation. Although GW3965 treatment induced a high expression of SREBF1 in comparison with control conditions, the treatment did not increase mDAN numbers and did not rescue the LBX1 KD effect (Figure 6F-G and Supplementary Figure 4). Thereby, suggesting that the critical role of LBX1 in mDAN differentiation lies upstream of cholesterol metabolism.

Further exploration of the DEGs altered by LBX1 KD highlighted the downregulation of eIF2 signaling and alterations in the regulation of eIF4 and p70S6K signaling (Figure 6B). Consistently, we found mTOR signaling to be predicted as downregulated upon LBX1 KD (mTOR signaling, -log(p-value) = 3.97, z-score = −0.707, Supplementary Table 4). It has been previously shown that mTOR signaling can control lipid metabolism through SREBF1, and mTOR inhibition leads to eIF4 sequestering by the eukaryotic initiation factor 4E-binding proteins (4E-BPs), impeding translation (90, 91). Therefore, downregulation of mTOR signaling due to the LBX1 KD could explain the observed downregulation in translation and cholesterol biosynthesis. Indeed, we found sirolimus (rapamycin), an mTOR inhibitor, as a chemical drug showing increased predicted activity among upstream regulators in our IPA analysis (activation z-score = 3.031, p-value of overlap = 1.21e-14, Supplementary Table 4). Therefore, the role of mTOR signaling during mDAN differentiation was tested using rapamycin. Neurons were treated with rapamycin as they were treated with GW3965, under normal differentiation and under LBX1 KD. Rapamycin treatment downregulates LBX1 in cells only transduced with shScramble (Figure 6F). In addition, rapamycin treatment was able to downregulate SREBF1 and SREBF2 in a similar way as observed before for LBX1 KD alone. Lastly, rapamycin treatment was also able to decrease the number of mDANs present in cultures (Figure 6G). However, rapamycin did not induce a similar loss of overall cell numbers as observed upon LBX1 KD (Supplementary Figure 4).

In summary, LBX1 KD results in an extensive deregulation of the neuronal transcriptome. Cholesterol biosynthesis appeared as one of the main affected pathways regulated by LBX1, possibly through the regulation of SREBF1/2 and NR1H3/2. Stimulating this pathway using GW3965 did not result in increased numbers of mDANs, suggesting that misregulation of the TFs involved in lipid metabolism alone is insufficient to explain the observed impact on mDANs. Perturbation of mTOR signaling provides a possible explanation of the results observed upon LBX1 KD. Indeed, mTOR inhibition was able to reproduce many of the changes induced by LBX1 KD.

## Discussion

Here we present a multi-omics data integration approach to discover novel TFs controlling mDAN differentiation. Although previous studies have used similar transcriptomic and epigenomic profiling of mDANs (16, 18), our assessment of human mDAN differentiation is cell-type-specific and time-resolved. Together with the *de novo* identification of key regulators of mDANs, we also provide a functional validation and characterization of the newly identified TFs.

The EPIC-DREM pipeline here offers an unbiased and data-driven tool for identifying TFs controlling processes across time based on transcriptomic and epigenomic data. The workflow can also be used for processes other than differentiation such as drug treatments and disease progression. This study used EPIC-DREM for the first time in conjunction with ATAC-seq data. Previously, EPIC-DREM operated on ChIP-seq datasets (8) and some of the tools integrated into this pipeline have been optimized for the use of other methods to detect accessible regions such as DNaseI-seq and NOMe-seq (22). EPIC-DREM can be applied to an ample range of epigenomic datasets that together with transcriptomic data can help to uncover key regulators and biological insights.

For the first time, we identified TFs controlled by SEs during mDAN differentiation. A total of 49 TFs were found to be under the control of SEs between day 30 and day 50 of differentiation (Supplementary Table 2). Of these, 17 were among the top TFs predicted by EPIC-DREM (Figure 3B). Among the remaining 32 TFs that did not overlap with EPIC-DREM, we found additional promising candidates. For example, ZFHX4 and CSRNP3 are highly expressed in iPSC-derived mDANs (data not shown) and in developing human mDANs *in vivo* (http://linnarssonlab.org/ventralmidbrain/, (13). These TFs were not selected in our analysis because their motifs are not known, namely, their TFBS in the accessible regions cannot be determined through footprinting and the consequent prediction as regulators was not possible. This highlights a limitation of the approach used here, but simultaneously opens new research avenues for previously less studied TFs that seem to play an important role in neurogenesis.

Gene perturbation experiments were performed to validate the identified candidates: HOXB2, LBX1, NHLH1, NR2F1, and NR2F2. After TF depletion during differentiation, it was found that LBX1, NHLH1 and NR2F1/2 presented a stronger phenotype than a well-established and characterized TF involved in mDAN neurogenesis: LMX1B. Further characterization of the role of these TFs revealed that only LBX1 and NHLH1 inductions during differentiation were able to increase mDAN numbers, while induction time was critical for LBX1.

Overall, our results are in line with previous findings. Loss of function in the LBX1 protein, specifically in the protein-protein interaction domain, in addition to being lethal during development due to cardiac abnormalities in mice, has been shown to impair neurogenesis during the development of the neural tube (92). This finding can be correlated with what was observed during LBX1 KD in mDANs, where differentiating cells always struggle to survive. Moreover, the findings on the development of spinal cord interneurons, RTN, and dorsal neurons from the hindbrain have linked LBX1’s function to cell fate decisions (67, 68, 92). Here we have added a role for LBX1 in mDAN differentiation.

On the other hand, NHLH1 loss of function has been shown to be lethal in mice, but only in adult life. However, if accompanied by loss of NHLH2, mice die already at birth (72, 93). Furthermore, NHLH1 has been shown to control neurogenesis by regulating the expression of neuronal-specific genes during *Xenopus* development (94). Our results support an important role for NHLH1 in controlling mDAN differentiation, consistent with previous findings.

After seeing the relevance of LBX1 and NHLH1 during differentiation, transcriptomic analysis after TF depletion was performed to determine the main processes controlled by these TFs. LBX1 KD showed a clear downregulation of cholesterol biosynthesis-related genes together with SREBF1 and SREBF2. As SREBF1 is at the same time regulated by LXRα/β, and all of them have been previously related to mDAN neurogenesis, it was decided to stimulate this pathway using GW3965. However, the treatment with this molecule was not sufficient to rescue the LBX1 KD effect or increase mDAN numbers in the differentiation protocol used for this study. It is possible that for GW3965 treatment to be effective, it should be either added from the beginning of differentiation or cells should be treated for a more extended period. We refrained from these experiments because the main effects of LBX1 KD were studied late in differentiation during the first 5 days of maturation.

On the other hand, as stimulation of TFs involved in cholesterol metabolism did not rescue the LBX1 KD effect, we focused on another enriched pathway, eIF signaling. This pathway is involved in translation, a process also downregulated by LBX1 KD. Exploration of the data for a common origin of cholesterol biosynthesis and translation downregulation revealed mTOR signaling inhibition as a possible explanation and in line with previous findings (90, 91, 95–97). mTOR signaling was recently shown to play different roles in mDAN biology, controlling their morphology, dopamine release, and electrophysiology (98). Indeed, inhibiting mTOR signaling using rapamycin recapitulated most of the features observed by LBX1 KD.

Transcriptional profiling and more detailed analysis of neurons upon NHLH1 KD revealed a downregulation of the mature form of miR-124-3p. This relationship between NHLH1 and miR-124 has not been described before. miR-124 is a potent neurogenic microRNA that has been found to be downregulated in PD patients (99) and shown to be protective in PD animal models (100, 101). After determining the capacity of ABX464 in inducing miR-124 levels in neurons, the molecule was used to determine its effect during mDAN differentiation and as an attempt to rescue the NHLH1 KD. It was found that for inducing at least a 2-fold increase of miR-124-3p, the concentration needed was causing neurotoxicity, possibly due to the already high expression of miR-124 in the neuronal lineage. Independently of that, ABX464 treatment was able to increase mDAN numbers and to induce NHLH1 expression. Therefore, NHLH1 and miR-124 seem to be part of a positive feedback loop. To unveil potential applications of ABX464, its neurotoxicity should be reduced for example by chemical modification of the molecule. ABX464 is already in clinical trials for the treatment of rheumatoid arthritis and ulcerative colitis due to its anti-inflammatory potential (87, 102). Namely, ABX464 safety and efficacy has been already evaluated and the repurposing of its use for PD could be taken into consideration to increase miR-124-3p levels as a neuroprotective treatment.

## Data availability

The source RNA-seq, ATAC-seq and ChIP-seq fastq files have been deposited at https://ega-archive.org/, under the accession number EGAD00001009288. Additional intermediate files such as RNA-seq counts, ATAC-seq peaks and bigwig files, and ChIP-seq H3K27ac SEs and bigwig files, can be provided upon request.

Analysis code can be found in the following repository: https://gitlab.lcsb.uni.lu/borja.gomezramos/gomezramos_et_al_2023/-/tree/main/

## Acknowledgments

The authors would like to thank Marie Catillon for support with experiments. The computational analysis presented in this paper were carried out using the HPC facilities of the University of Luxembourg.

## Author Contributions

B.G.R., J.O., M.S., T.S., R.K. and L.S. conceived the project and designed the experiments. B.G.R. performed most of the experiments and bioinformatic analysis. J.O. established and performed ATAC-seq protocol and performed bioinformatic analysis. N.d.L., Ro.K., and M.S. established the EPIC-DREM pipeline and performed the analysis. A.G. designed the RNA-seq pipeline and performed bioinformatic analysis. A.R. designed and generated the iPSC reporter line in consultation with C.K.. R.H. prepared RNA-seq libraries and performed the sequencing. F.M. performed rapamycin treatments. Ro.K., M.S., T.S., R.K., and L.S. provided input on data interpretation and revision of the study. L.S. supervised the project. B.G.R. and L.S. wrote the manuscript. All authors revised and approved the final manuscript.

## Supplementary Information

The Supplementary Information contains four supplementary figures and four supplementary tables.

