## Supplementary Information for "Multi-omics analysis identifies LBX1 and NHLH1 as central regulators of human midbrain dopaminergic neuron differentiation"

#### Supplementary Figures

**Supplementary Figure 1.** Heatmap showing the expression of cell-type specific markers of smNPCs, neurons, mDANs, and mDAN subtypes selected from literature (13, 49).

**Supplementary Figure 2. A)** Live imaging for early and late transductions with shRNA lentiviral particles across time. Bar plots representing GFP+ cells at day 15 of analysis **B)** Live imaging for early and late transduction with lentiviral particles for the overexpression of LBX1, NHLH1 and NR2F2. Bar plots representing GFP+ cells at day 15 of analysis. Scale bar = 200 µm

**Supplementary Figure 3. A)** Loci and expression of the three genes coding for miR-124 **B)** Live imaging of cells under ABX464 treatment and NHLH1 KD at day 14 of differentiation **C)** Flow cytometry analysis showing the differences in shape between neurons treated with ABX464 and non-treated **D)** Expression Ct values detected by TaqMan and RT-qPCR for miR-1243p and NHLH1, respectively, in samples treated with ABX464 and under NHLH1 KD.

**Supplementary Figure 4.** Live imaging of cells under GW3965 and Rapamycin treatment and LBX1 KD at day 14 of differentiation. Scale bar = 200 µm.

Supplementary Figure 1

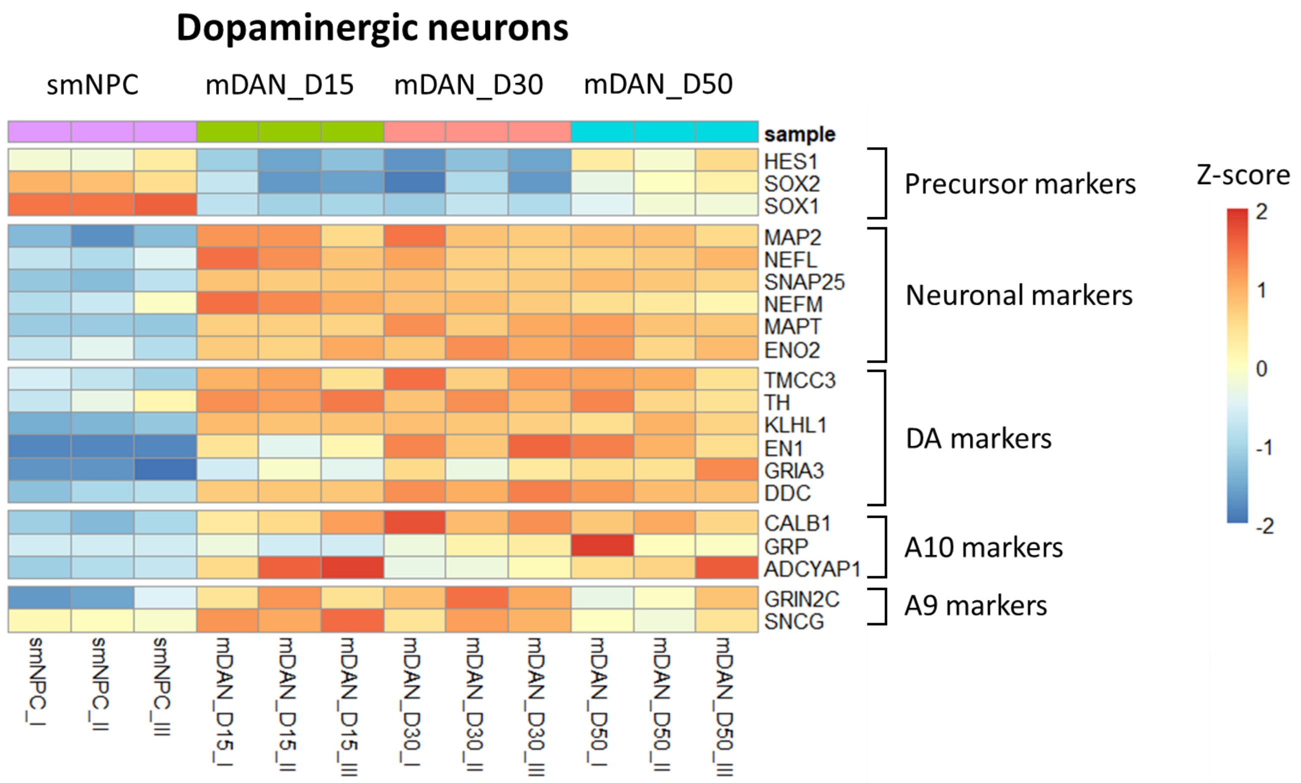

Supplementary Figure 2

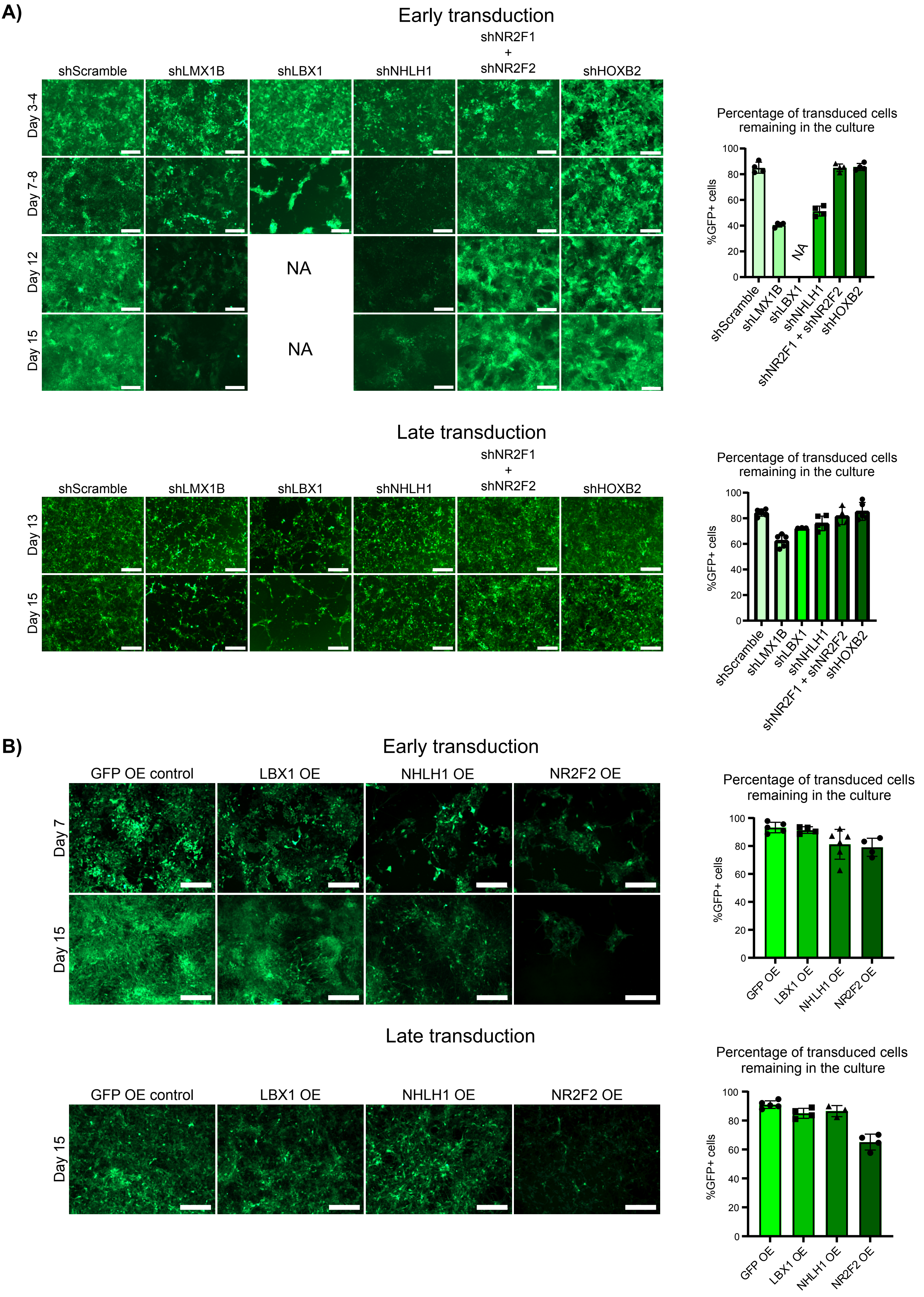

### Supplementary Figure 3

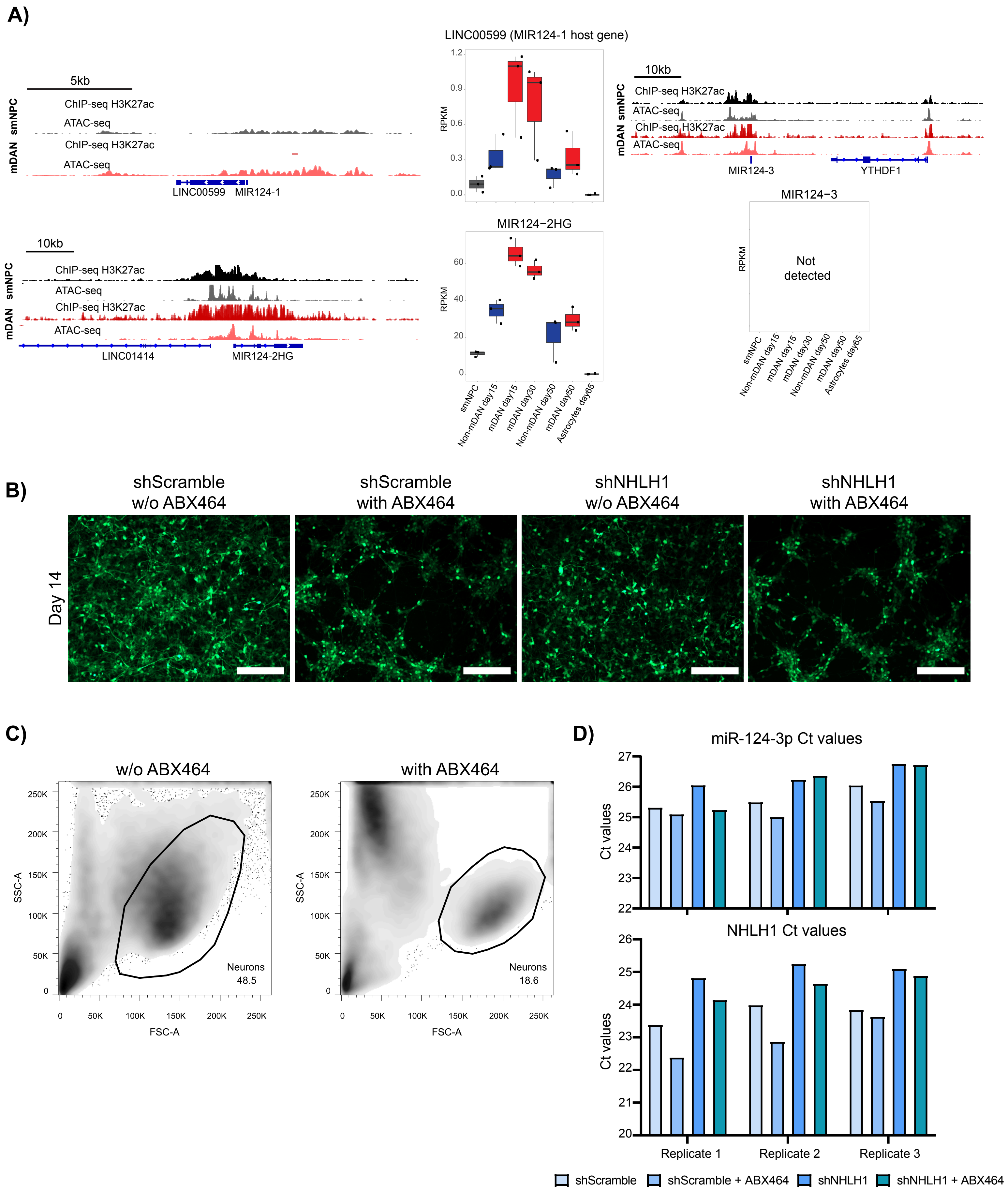

Supplementary Figure 4

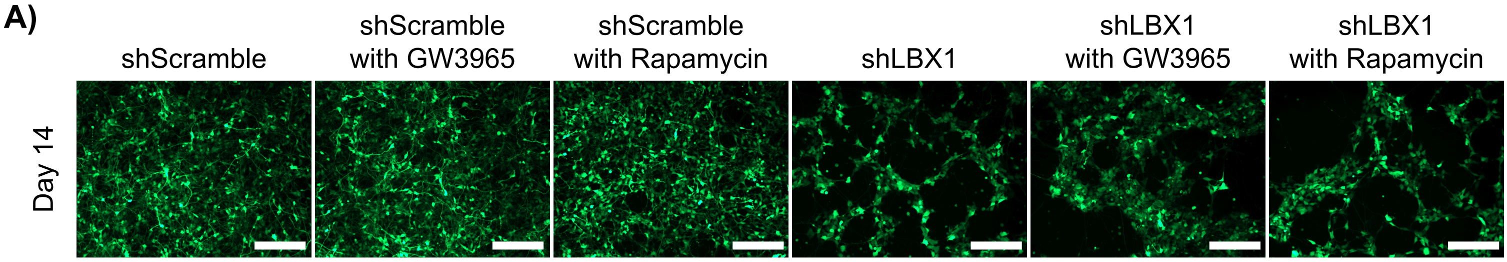

#### Supplementary Tables

**Supplementary Table 1.** EPIC-DREM ranking of TFs per split node.

**Supplementary Table 2.** List of TFs controlled by SEs at days 30 and 50 in mDANs. List of all TFs controlled by SEs in mDANs. List of top 20 TFs across all nodes from EPIC-DREM. Common TFs after overlapping the list of all TFs controlled by SEs in mDANs with the list of top 20 TFs across all nodes from EPIC-DREM.

**Supplementary Table 3.** List of DEG from late transduction NHLH1 KD RNA-seq with a padj < 0.05. IPA analysis from NHLH1 KD RNA-seq data using the list of DEG. List of predicted miR-124-3p targets from TargetScan 8.0 using the list miR-124-3p.1 and filtering for the genes expressed in the RNA-seq data from NHLH1 KD. Common targets after overlapping the list of upregulated genes from NHLH1 KD RNA-seq DEG list and the predicted expressed targets of miR-124-3p from TargetScan.

**Supplementary Table 4.** List of DEG from late transduction LBX1 KD RNA-seq with a padj < 0.05. IPA analysis from LBX1 KD RNA-seq data using the list of DEG, showing pathway enrichment analysis and predicted upstream regulators
